# Multivariate Coevolution Shapes Life-History Strategies Across Amniotes

**DOI:** 10.1101/2025.03.09.642183

**Authors:** Yang Zhang, Shan Huang, Muyang Lu, Yunshu Zhang, Qin Li, Fangliang He, Chunhui Hao

**Affiliations:** School of Life Sciences, Nanjing University, Nanjing, 210023, China; School of Geography, Earth and Environmental Sciences, University of Birmingham, Edgbaston, Birmingham, B15 2TT, UK; College of Life Science, Sichuan University, Chengdu, 610064, China; ECNU-Alberta Joint Lab for Biodiversity Study, School of Ecological and Environmental Sciences, East China Normal University, Shanghai, 200241, China; Department of Renewable Resources, University of Alberta, Edmonton, AB, T6G 2H1, Canada

**Author notes:** Corresponding author: Chunhui Hao, School of Ecological and Environmental Sciences, East China Normal University, No. 500, Dongchuan Road, Shanghai, 200241, China. These authors contributed equally (Yang Zhang, Chunhui Hao). **Statement of authorship** Study design: C.H.; Data collection: Y.Z. and Y.S.Z.; Methodology: Y.Z. and C.H.; Analysis: Y.Z. and C.H.; Visualization: Y.Z. and C.H.; Funding acquisition: Y.Z.; Supervision: S.H., M.L. and F.H.; Writing—original draft: Y.Z. and C.H.; Writing— review and editing: Y.Z., S.H., M.L., Q.L., F.H. and C.H. **Data accessibility statement** All data generated or analyzed in this study are included in the Supplementary Materials. Code will be publicly archived upon publication in GitHub repositories.

**Keywords:** life-history, Amniotes, coevolution, phylogenetic comparative analysis, phylogenetic path analysis, multivariate Ornstein–Uhlenbeck model

## Abstract

Life-history traits jointly determine survival, growth, and reproduction, yet their evolutionary interdependence remains poorly understood. Most comparative studies focus on pairwise trade-offs or composite axes, overlooking the complexity of multivariate coevolution. Here, we integrate phylogenetic path analysis and multivariate Ornstein–Uhlenbeck models to investigate the coevolution of five key life-history traits across amniotes. We show that life-history evolution proceeds through modular coevolutionary pathways with distinct tempos among traits and taxa. Despite loading similarly on the fast–slow axis of mammals and birds, clutch size and clutch frequency evolve independently at both instantaneous and long-term timescales, with clutch frequency adapting rapidly, whereas clutch size evolves slowly over macroevolutionary time. Lifespan, development time, and offspring size exhibit positive coevolution at instantaneous scales across clades, but follow distinct adaptive modes with varying directions over longer timescales. These patterns become evident only through the integrated multivariate comparative approaches.

## Introduction

Life-history traits collectively determine the survival, growth, and reproductive success of organisms. Their covariation across taxa reflects how species allocate limited resources such as energy, space, and time to maintain fitness (De Jong 1993; Stearns 1992). Theory predicts that these traits rarely evolve independently. From a genetic perspective, mechanisms such as pleiotropy, epistasis, and linkage disequilibrium couple traits, meaning that evolutionary change in one trait typically alters the selective pressures on other traits (Pavličev & Cheverud 2015; Saltz *et al*. 2017; Svensson *et al*. 2021). Similarly, physiological mechanisms such as endocrine regulation coordinate multiple life-history functions—including growth, fertility, immunity and survival— through shared hormonal pathways (Ketterson *et al*. 2009; Vitousek & Schoenle 2019). These genetic and physiological processes underscore that life-history evolution is inherently multivariate, where suites of traits coevolve as integrated networks.

Yet most comparative studies simplify this complexity. Traditional studies have primarily focused on pairwise relationships, such as the trade-off between offspring size and number (MacArthur and Wilson 1967; Pianka 1970; Stearns 1989), but such a bivariate approach is handicapped in dissecting the complexity of multi-trait coevolution and overlooks the potential of other traits in mediating apparent pairwise correlations (Stott *et al*. 2024; Thorson *et al*. 2023). As a result, a pairwise trade-off can appear weak, absent, or even reversed in some lineages if the effect of other traits is not taken into account. For example, the expected negative correlation between offspring size and number becomes weak or even disappears in some insect lineages when adult body size is not considered (Bakewell *et al*. 2020; Flatt 2020).

As an attempt to synthesize the multidimensionality in life-history evolution beyond bivariate analyses, principal component analysis (PCA) is often employed to identify major axes of trait covariation such as the fast–slow continuum (Sæther 1987; Stearns 1983). However, reliance on the fast–slow axis can obscure additional, and often biologically meaningful covariation in life-history traits. For example, mammals consistently exhibit a second, independent precocial–altricial axis beyond the fast–slow model (Del Giudice 2020; Dobson & Oli 2007; Stearns 1983). Likewise, marine turtles occupying the ‘slow’ end of the fast–slow axis combine long lifespans, delayed maturation, and high survival with surprisingly high reproductive output—contrary to classic fast–slow predictions that expect reproduction to trade off against survival (Stott *et al*. 2024; Wright *et al*. 2020). Moreover, these composite principal axes (as a linear combination of individual traits) are often difficult to interpret; they do not allow explicit inference of the tempo and mode of trait coevolution, or testing hypotheses regarding the coevolutionary processes underlying multivariate patterns (Del Giudice 2020; Stott *et al*. 2024; Svensson *et al*. 2021; Walsh & Blows 2009).

In this study, we directly examined the multivariate interaction of life-history traits and further, modelled their coevolution using an integrative multivariate phylogenetic framework that combines phylogenetic path analysis (PPA) and multivariate Ornstein– Uhlenbeck (OU) models. We tested whether life-history traits interact and coevolve, and how, across three major amniote clades: mammals, birds, and reptiles. Amniotes constitute one of the most diverse vertebrate lineages and display substantial variation in reproductive strategies and life-history schedules, making them ideal systems for understanding macroevolutionary patterns (Jeschke & Kokko 2009; Myhrvold *et al*. 2015). Our results showed that life-history strategies are governed by distinct, modular coevolutionary pathways, with the tempo and direction of evolution varying across clades — insights that were only revealed by the integrated multivariate comparative approaches.

## Materials and methods

### Data collection

We compiled species-level life-history trait data for mammals, birds, and reptiles using the Amniote Database (Myhrvold *et al*. 2015) as the primary source. Additional datasets from Allen *et al*. (2017), Bakewell *et al*. (2020) and Capellini *et al*. (2015) were integrated, with newer records prioritized to resolve overlaps, supplement missing traits and increase species coverage.

To ensure robust inference in fitting multivariate Ornstein-Uhlenbeck (OU) models (Cooper *et al*. 2016), we retained only species with complete data for five key life-history traits: offspring size (OS, in grams), development time (DT, time from independence to sexual maturity in days), adult reproductive lifespan (AL, in days), clutch size (CS, number of offspring or egg per reproductive event), and clutch frequency (CF, number of litters or clutches per year). These traits are fundamental determinants of population growth dynamics in a given species (Sæther *et al*. 2013) and have been widely used in comparative studies of life-history strategies (Allen *et al*. 2017; Morrow *et al*. 2020). Definitions and measurement details were provided in Supplementary Methods (section 1). The final dataset comprised 1341 amniote species, including 774 mammals, 156 birds, and 411 reptiles (Table S1). All trait values were natural log-transformed to meet the assumptions of the statistical methods.

Additionally, we controlled for the influence of adult body size, which is a known key allometric determinant shaping most life-history traits and metabolic rates (Beccari *et al*. 2024; Hallmann & Griebeler 2018; Jeschke & Kokko 2009). We calculated the body size-corrected residuals for each life-history trait (Supplementary Methods section 2), and all main analyses were conducted using these size-adjusted traits.

Phylogenetic trees used in this study were obtained and pruned from the VertLife Database (https://vertlife.org/; accessed 25 June 2025), following procedures described in the Supplementary Methods section 3.

### Baseline multivariate pattern derived from principal components analysis

To establish baseline expectations for comparison with other multivariate approaches (see below), we used phylogenetically-informed principal components analysis (pPCA) to examine the covariation among life-history traits in our dataset and assessed its alignment with the fast–slow continuum (Charnov 2002; Sæther 1987; Stearns 1992), while accounting for the phylogenetic non-independence in trait covariation (Jeschke & Kokko 2009; Revell 2009). We defined the fast–slow continuum using the first principal component (PC1), consistent with the previous definition (Beccari *et al*. 2024; Jeschke & Kokko 2009). This analysis was performed using the *phyl*.*pca* function from the R package “*phytools*” (version 2.4.4; Revell 2024).

### Multivariate interaction from phylogenetic path analyses

We employed phylogenetic path analysis (PPA) to explicitly characterize the multivariate relationships—both direct and indirect—among individual life-history traits. PPA extends the structural equation modelling (SEM) approach by evaluating the goodness-of-fit of data to hypotheses concerning the relationships among multiple variables while accounting for phylogenetic non-independence (Gonzalez-Voyer & Von Hardenberg 2014; Hardenberg & Gonzalez-Voyer 2013). Similar to SEM, PPA uses the d-separation method to test conditional independencies predicted by competing models (Shipley 2000, 2013), enabling a comprehensive assessment of the interconnected pathways underlying life-history evolution across species.

We formulated 32 candidate models, each represented as a directed acyclic graph (DAG), to decipher the hypothesized multivariate associations among five life-history traits (Figure S9), using the equation syntax detailed in Table S4. We formulated hypothetical models following these rules: (i) all consistent pairwise associations identified across all taxa via PGLS were incorporated as fixed paths in every candidate model (Figure S9; Table S4); (ii) for associations that varied across taxa, we systematically adjusted their inclusion across different models, thereby generating a set of alternative hypothesis models for comparative evaluation (Figure S9; Table S4).

We conducted PPA using the R package “*phylopath*” (version 1.3.0; Van Der Bijl 2018), under an Ornstein–Uhlenbeck (OU) model of trait evolution, which was selected based on its fit to the evolutionary dynamics of these traits across taxa (Figures S2-S4). Goodness-of-fit was assessed via Fisher’s C statistic, where a model was considered a good fit if the C statistic was non-significant (*P* > 0.05) (Shipley 2000, 2013). Moreover, to rank the candidate models and identify the best-supported model, we calculated the C statistic information criterion (CICc) for each model. The best-supported model was the one with a non-significant C statistic (*P* > 0.05) and the lowest CICc value. Models with ΔCICc < 2 relative to the best model were considered statistically equivalent; we thus computed model-averaged estimates from all top models to obtain an averaged best model (Shipley 2013).

To understand how multivariate patterns from PPA modify the bivariate correlations typically observed, we employed phylogenetic generalized least squares (PGLS) to identify pairwise relationships among all size-corrected life-history traits across species within each taxon, accounting for phylogenetic non-independence. PGLS can incorporate various evolutionary models (e.g., Pagel’s λ, Brownian motion, or early burst) to define the distribution of the residuals from the model (Freckleton *et al*. 2002). For each pair of size-corrected life-history traits, we performed PGLS using the best-supported evolutionary model selected by the lowest AIC value. The PGLS analyses were carried out using the R package “*phylolm*” (version 2.6.5; Tung Ho & Ané 2014). To ensure robust parameter estimation, we conducted 1,000 independent bootstrap replicates for each PGLS model. Additionally, we derived regression coefficients along with their associated statistical outputs, including 95% confidence intervals, *P*-values, and *R*^*2*^ statistics, to assess the goodness-of-fit of each model (Figures S5-S7; Table S2).

### Modelling the multivariate coevolution among traits using multivariate Ornstein– Uhlenbeck (OU)-based models

PPA provides a clear picture of the multivariate associations among life-history traits, but does not explicitly capture the evolutionary processes that generate these associations. To extend beyond correlational patterns and explicitly model how traits coevolve along the phylogeny, we complemented PPA with multivariate Ornstein– Uhlenbeck (OU) models (Gonzalez-Voyer & Von Hardenberg 2014; Van Der Bijl 2018). The multivariate OU framework enables us to test the relative likelihood among alternative causal directions (i.e., A → B or B → A) of trait coevolution, and to quantify both the instantaneous and long-term modes of coevolutionary dynamics (Bartoszek *et al*. 2023, 2024).

Speciafically, the multivariate OU process extends the univariate OU process (Butler & King 2004; Hansen 1997), which describes adaptive evolution of a trait toward an optimal value. This framework models the joint evolution of a *k*-dimensional suite of traits, 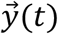, over a period of time through the following stochastic differential equation:

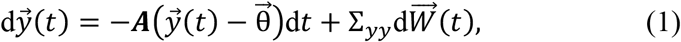

Where 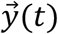 is the vector of trait values at time *t*; ***A*** matrix represents rates of adaptation of trait values towards optimum values. Each off-diagonal entry in the ***A*** matrix represents the instantaneous rate of co-adaptation between traits, describing how a small deviation in one trait affects the expected rate of change in another at a given moment in the stochastic process; 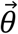 is the vector representing the optimum values for these traits; 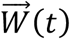 is a *k*-dimensional standard Brownian motion. The diffusion matrix Σ_*yy*_ scales the resulting stochastic perturbations, which represent non-directed changes in trait values attributable to unconsidered selective factors, environmental fluctuations, or evolutionary constraints (Bartoszek *et al*. 2012, 2023, 2024; Clavel *et al*. 2015). Model fitting was performed with the R package “*mvSLOUCH*” (2.7.6; Bartoszek *et al*. 2024).

In the initial step, we parameterized the ***A*** matrix to reflect the relationships specified in each of the 32 DAGs defined previously (Figure S9). Each ***A*** matrix thus corresponded to a distinct hypothesis of coevolutionary structure among traits, and we compared the relative likelihoods of these competing models (Figure M1; see Supplementary Methods section 6 for details). To ensure robust parameter estimates and model selection, each model configuration was run 500 times with different starting points of optimization. The best-supported model for each animal group was identified as the one with the lowest AICc value across all runs.

In the second step, we further examined the causal structures of the best-supported model identified in the first step by evaluating the relative likelihood of alternative directional hypotheses for each coevolutionary link. Each possible direction (e.g., A → B vs. B → A) represents a distinct hypothesis describing how deviation in one trait affects the expected rate of change of another trait toward (or away from) its adaptive optimum within the OU framework (Reitan *et al*. 2012). In the multivariate OU model, such directionality is reflected in the asymmetry of the off-diagonal entries in the ***A*** matrix (Figure M1; see Supplementary Methods section 6 for details).

The ***A*** matrix of the final causal model—selected based on the smallest AICc—was then decomposed to obtain its eigenvalues and eigenvectors, allowing inference of long-term evolutionary dynamics of traits. The eigenvalues (λ) of ***A*** describe the relative long-term rates at which linear combinations of traits evolve toward their optima along the independent directions in trait space defined by the corresponding eigenvectors 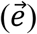 of ***A*** (Bartoszek *et al*. 2023, 2024). We also reported the evolutionary half-life 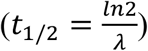 for each eigenvector, which represents the time required for an eigenvector to evolve halfway from its ancestral state to the optimum (Hansen & Bartoszek 2012). For comparative clarity, these half-lives were additionally expressed as a relative percentage of total tree height (Figure M1; see Supplementary Methods section 6 for details).

## Results

### Baseline patterns revealed by principal components analysis

Our pPCA analyses show that the first principal component (PC1) captured a more consistent fast–slow continuum in mammals and birds but not in reptiles, underscoring both the generality and limitations of this classic framework (Bakewell *et al*. 2020; Jeschke & Kokko 2009; Salguero-Gómez *et al*. 2016). Specifically, in both mammals and birds, PC1 reflected that species at the “fast” end of the continuum exhibited shorter lifespans, earlier maturation, and larger, more frequent clutches, while “slow” species displayed the opposite patterns (Figures 2a and 2b). In contrast, reptiles showed a reversed reproductive strategy: PC1 primarily represented the trade-off between clutch frequency and other traits, suggesting that fast-strategy species produced smaller, but more frequent clutches, whereas slow-strategy species had larger, less frequent ones (Figure 2c).

**Figure 1.**
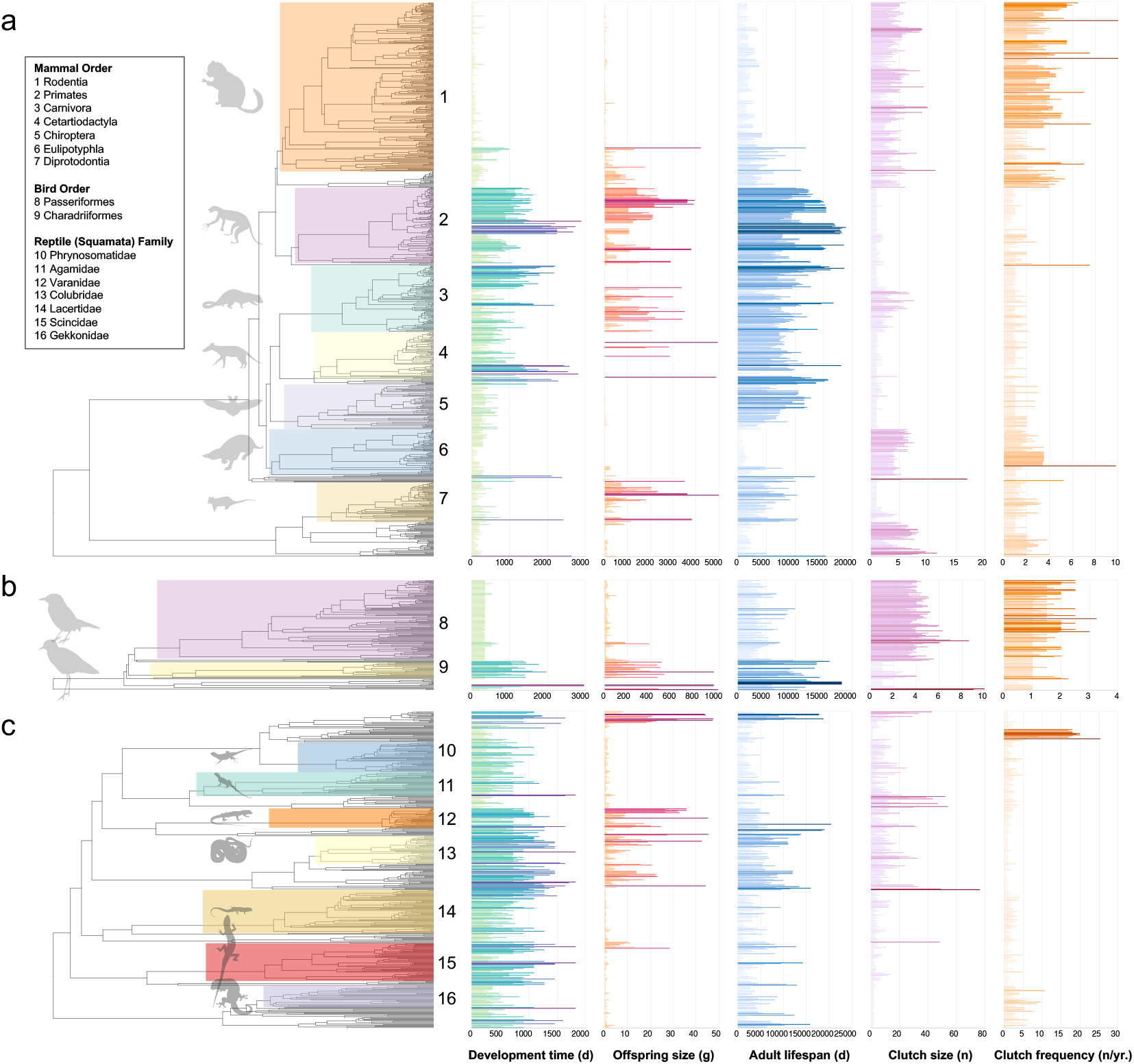
Phylogenetic relationships and associated life-history trait values for (a) 774 mammalian, (b) 156 avian, and (c) 411 reptilian species. The phylogeny of clade Amniota is displayed with numbered clade labels corresponding to the major orders and families indicated at the plot peripheries. Life-history traits, including offspring size, clutch size, clutch frequency, development time, and lifespan, are mapped onto the trees to illustrate their variation across lineages. Bar length corresponds to trait value. Extreme trait values falling outside the bounds of the x-axis were excluded to improve visualization clarity. The complete dataset is available in Supplementary Materials (Table S1).

**Figure 2.**
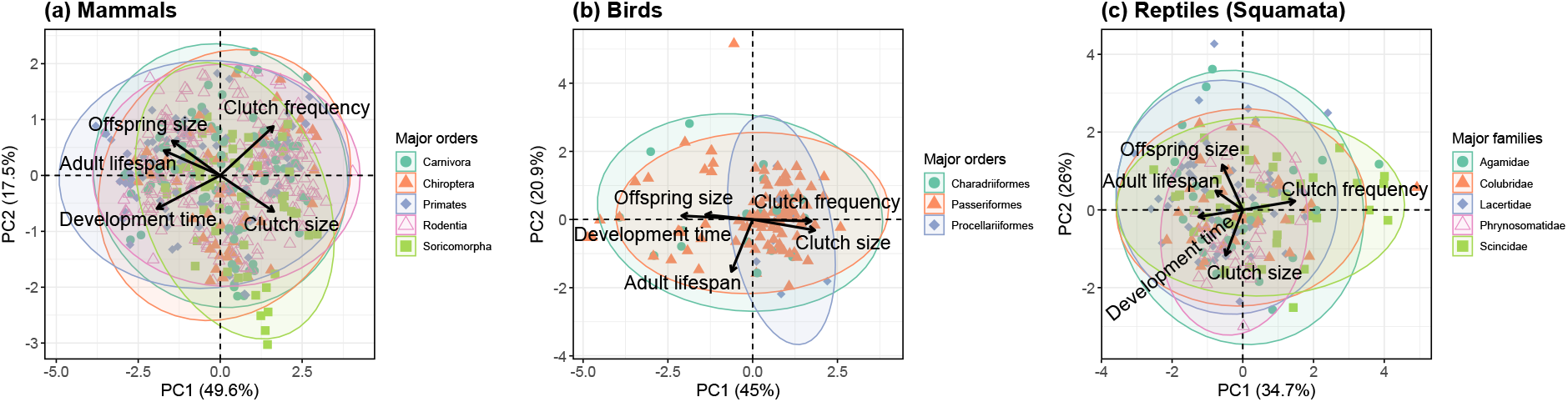
Biplots of phylogenetically informed PCA illustrating the covariations among five life-history traits in (a) 774 mammalian, (b) 156 avian, and (c) 411 reptilian (squamate) species. Arrows represent the loadings of five key traits on the first two principal components (PCs). Across all groups, PC1 primarily captures the fast–slow continuum of life-history strategies, while PC2 reflects secondary dimensions of trait covariation. Each point represents a species, with point shapes indicating major clades (five mammalian orders, three avian orders, and five squamate families) and colors corresponding to these clades within each group. Ellipses summarize the distribution of each clade in multivariate trait space, emphasizing both shared patterns and clade-specific divergences. Collectively, the first two axes explain 67.1% of the total variation in mammals, 65.9% in birds, and 60.7% in reptiles.

### Distinguishing direct and indirect associations among life-history traits

Within the PGLS framework, we found that multivariate models consistently provided a better fit than univariate models, indicating that the evolution of life-history traits is better explained by multivariate associations rather than by simple bivariate relationships (Figure 3b; Table S3; see Supplementary Method section 4 for details).

**Figure 3.**
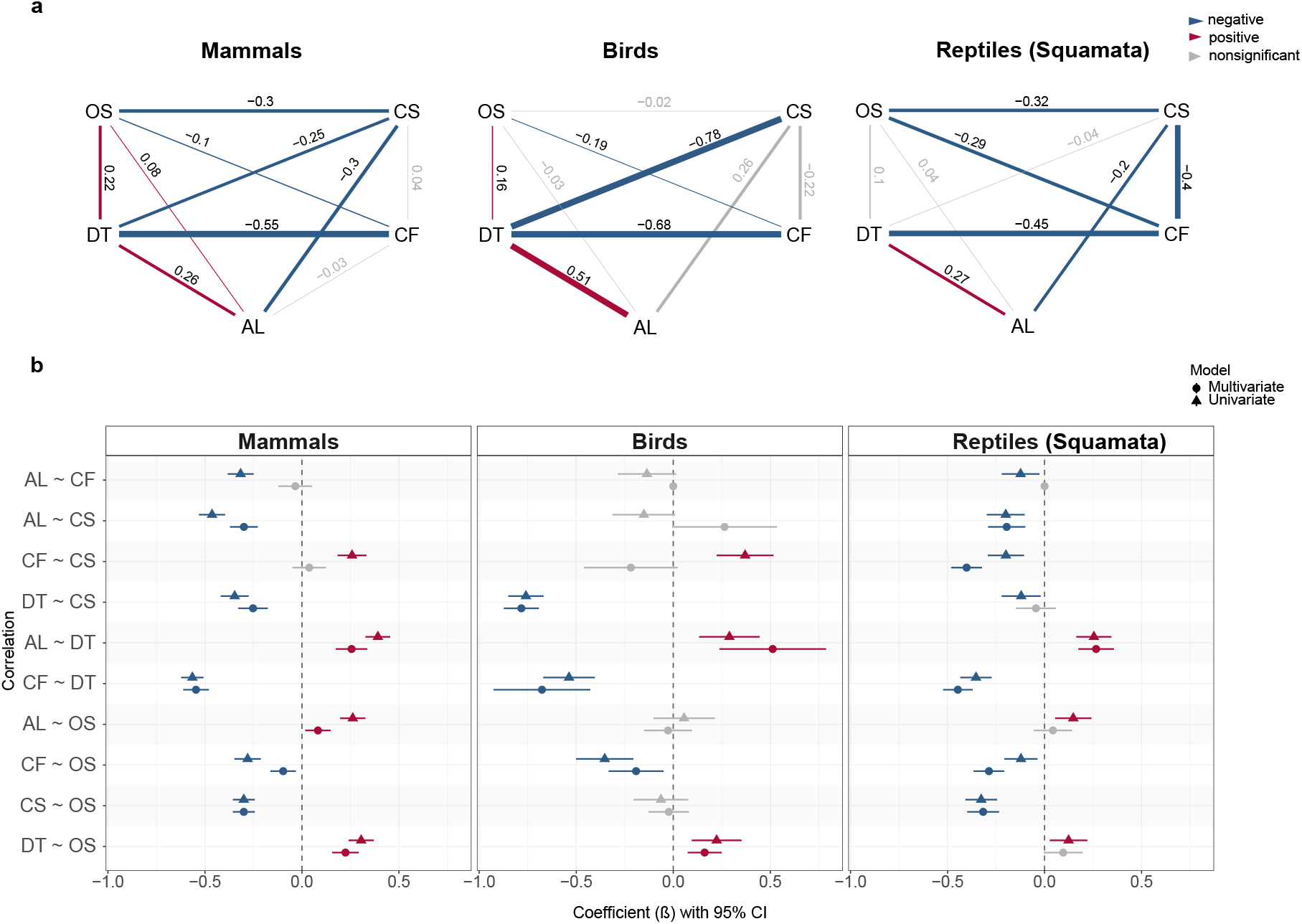
(a) Averaged best models of phylogenetic path analysis (PPA) for mammals, birds, and reptiles. Red and blue lines represent significant positive and negative associations between pairs of traits, respectively, with the thickness of each line reflecting the magnitude of their standardized path coefficients. Gray lines indicate non-significant relationships between traits. The values adjacent to the lines denote the standardized average path coefficients. (b) Comparison of standardized regression coefficients from univariate (PGLS) and multivariate (PPA) models. Forest plots show regression coefficients with ± 95% confidence intervals for each pair of traits. Red and blue circles or triangles represent significant positive and negative associations, respectively, while gray circles or triangles indicate non-significant relationships. Triangular markers represent the coefficients derived from the PPA, whereas circular markers correspond to those from the PGLS models. Trait abbreviations: OS = offspring size; DT = development time; AL = adult lifespan; CS = clutch size; CF = clutch frequency.

In the best PPA models, we found that some associations detected by PGLS were mediated by third variables (Figure 3; Table S6). For example, in mammals, the positive correlation between clutch size and clutch frequency, as well as the negative correlation between clutch frequency and lifespan, both identified in PGLS, was mediated by offspring size and/or development time in the PPA models (Figure 3; Table S6). Similar mediation patterns were also observed in birds and reptiles (Figure 3; Table S6).

Moreover, PPA recovered patterns of life-history covariation across amniotes, consistent with the pPCA results (Figures 2 and 3a). Building on such consistency, PPA provided additional resolution by distinguishing direct from indirect associations among traits, thereby refining some of the relationships suggested by pPCA. For instance, while pPCA implied a possible positive correlation between clutch size and clutch frequency in mammals and birds (Figures 2a and 2b), PPA revealed that these associations were not direct but mediated through development time and offspring size (Figure 3a).

### Multivariate coevolution of life-history traits over the instantaneous timescale

We fitted multivariate OU models to explicitly test causal hypotheses regarding the coevolutionary coupling of life-history traits (Bartoszek *et al*. 2023, 2024). We found that for each group, models with diagonal Σ_*yy*_ consistently yielded lower AICc values (Wilcoxon test: *P* < 0.001; Figure 4a), indicating independent stochastic perturbations (see Supplementary Methods section 6 for details). Therefore, our subsequent analyses assumed independent stochastic perturbations.

**Figure 4.**
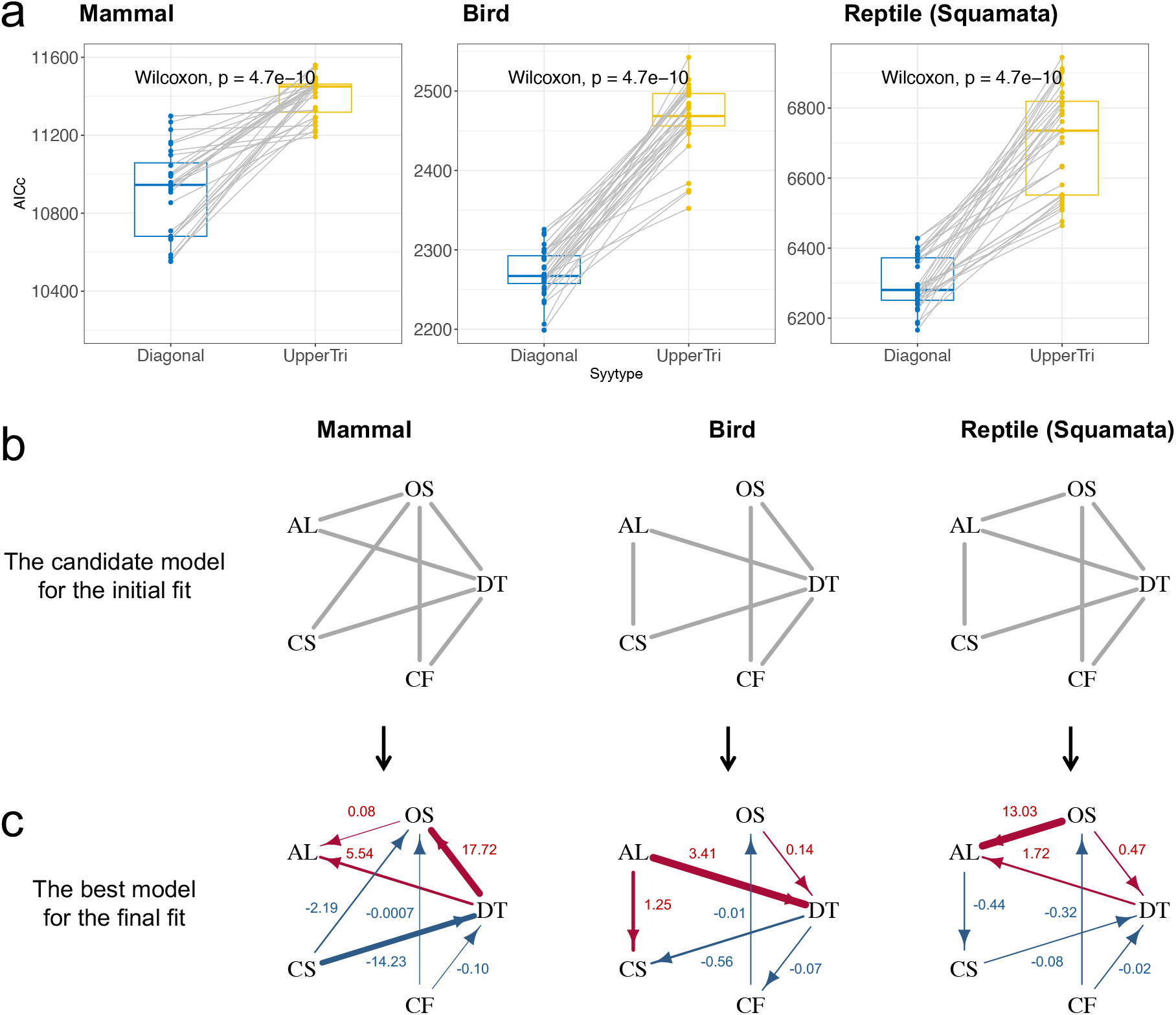
Multivariate Ornstein–Uhlenbeck (OU) models revealed instantaneous coevolutionary coupling among five life-history traits. (a) Model selection for the mode of stochastic perturbations. Dots represent the AICc scores for candidate models with either a diagonal or upper triangular Σ_*yy*_ matrix (the stochastic perturbation matrix). Across all amniote clades, models with diagonal Σ_*yy*_ consistently yielded lower AICc values, indicating that these life-history traits evolve primarily under independent stochastic perturbations rather than correlated selection. (b) The optimal model structure, selected from 32 candidate models (Figure S9), is shown for each major amniote group. (c) The causal model of trait coupling identified by the best-fitting multivariate OU model for each clade. Arrows represent coevolutionary links where a change in trait *j* drives evolutionary change in trait *i*. Red arrows indicate pull towards an optimal value, while blue arrows indicate push away from an optimum. Coefficients on arrows represent the strength of the coevolutionary coupling. Trait abbreviations: OS = offspring size; DT = development time; AL = adult lifespan; CS = clutch size; CF = clutch frequency.

The best-fitting OU model (i.e., with the lowest AICc) was a subset of the corresponding best PPA model in each amniote clade, differing only in the absence of certain direct coevolution parameters within the ***A*** matrix (Figure 4b, Table S8). Specifically, the OU models for all clades lacked a parameter for direct coevolution between clutch size and clutch frequency. Additionally, the best model for birds lacked a parameter between clutch size and adult lifespan, while the best model for reptiles lacked one between clutch size and offspring size (Figure 4b).

Across amniote groups, instantaneous coevolutionary relationships among life-history traits revealed both shared and lineage-specific patterns. Specifically, we found consistent positive instantaneous coevolution among development time, adult lifespan, and offspring size across groups, though direct coevolution between adult lifespan and offspring size was absent in birds (Figure 4c). In contrast, although clutch size and clutch frequency loaded similarly on the fast–slow axis in mammals and birds, no direct coevolution was detected, indicating their independent adaptation at an instantaneous timescale (Figure 4c). Moreover, the directions of arrows in the coevolutionary models varied across groups, which may reflect asymmetric, lineage-specific evolutionary histories of these species. For instance, in mammals and reptiles, deviations in development time influenced the expected rate of change in adult lifespan, whereas the opposite pattern was inferred in birds (Figure 4c).

Clutch size and frequency generally showed negative coevolution (trade-offs) with the other three traits in mammals and reptiles (Figure 4c). This aligns with trade-offs on the fast–slow axis in mammals but contrasts with that of reptiles, where clutch size loaded similarly to adult lifespan and development time on the fast–slow axis of reptiles (Figures 2a and 2c). In birds, adult lifespan and clutch size coevolved positively (Figure 4c) despite trade-offs on the fast–slow axis (Figure 2b). However, the two traits loaded similarly on the secondary axis (PC2) of birds, suggesting that their coevolution may reflect patterns associated with the secondary life-history strategy axis rather than the fast–slow continuum (Figure 2).

### Eigen-decomposition of the *A* matrix revealed complementary long-term modes of trait coevolution

Beyond instantaneous dynamics, long-term evolutionary patterns were revealed by the eigen-decomposition of the ***A*** matrix (Table 1). Across all clades, clutch frequency dominates the fastest evolutionary mode 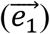 whereas clutch size dominates the slowest 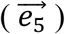, indicating that the pace of reproduction evolves independently from the investment per reproductive event (Table 1). This finding is robust to alternative datasets for birds with phylogenetically imputed trait values and datasets excluding offspring size (Figure S13; see Supplementary Methods section 6 for details).

**Table 1.** Eigen-decomposition of the ***A*** matrix in each amniote clade. Each eigenvector represents an independent axis of adaptation in the multi-trait space, with its corresponding eigenvalue (λ) determining the rate of adaptation toward the optimum. The rate of adaptation is then summarized by the evolutionary half-life (*ln*(2)/λ) and its proportion of the total tree height. Trait loadings for each eigenvector are displayed. For a given eigenvector, the trait that dominates the adaptive mode is highlighted in bold.

| a Mammal |  |  |  |  |  |
| --- | --- | --- | --- | --- | --- |
| Trait / Metric | $\rightarrow e_1$ | $\rightarrow e_2$ | $\rightarrow e_3$ | $\rightarrow e_4$ | $\rightarrow e_5$ |
| DT | 0.03 | 0.00 | 0.00 | 0.69 | 0.00 |
| OS | 0.00 | 0.00 | **1.00** | -0.53 | 0.00 |
| AL | 0.01 | **1.00** | 0.00 | -0.32 | 0.03 |
| CS | 0.00 | 0.00 | 0.00 | -0.38 | **1.00** |
| CF | **-1.00** | 0.00 | 0.01 | 0.00 | 0.00 |
| Eigenvalue ($\lambda$) | 104.55 | 65.03 | 45.36 | 42.33 | 30.63 |
| Half-life ($t_{1/2} = \frac{\ln 2}{\lambda}$) | 0.01 | 0.01 | 0.02 | 0.02 | 0.02 |
| Tree Height (%) | 0.66 | 1.07 | 1.53 | 1.64 | 2.26 |

| b Bird |  |  |  |  |  |
| --- | --- | --- | --- | --- | --- |
| Trait / Metric | $\rightarrow e_1$ | $\rightarrow e_2$ | $\rightarrow e_3$ | $\rightarrow e_4$ | $\rightarrow e_5$ |
| DT | -0.01 | 0.37 | 0.00 | 0.84 | 0.01 |
| OS | -0.01 | 0.09 | **1.00** | -0.54 | 0.00 |
| AL | 0.00 | 0.93 | 0.00 | 0.00 | 0.00 |
| CS | 0.00 | 0.00 | 0.00 | 0.00 | **1.00** |
| CF | **1.00** | 0.00 | 0.00 | 0.00 | 0.00 |
| Eigenvalue ($\lambda$) | 12.50 | 7.19 | 2.43 | 0.62 | 0.00 |
| Half-life ($t_{1/2} = \frac{\ln 2}{\lambda}$) | 0.06 | 0.10 | 0.29 | 1.12 | 147.53 |
| Tree Height (%) | 5.55 | 9.64 | 28.58 | 112.00 | 14752.96 |

**Table 1.** Eigen-decomposition of the $\mathbf{A}$ matrix in each amniote clade.
| c Reptile (Squamata) |  |  |  |  |  |
| --- | --- | --- | --- | --- | --- |
| Trait / Metric | $\rightarrow e_1$ | $\rightarrow e_2$ | $\rightarrow e_3$ | $\rightarrow e_4$ | $\rightarrow e_5$ |
| DT | -0.01 | 0.00 | 0.66 | -0.10 | -0.01 |
| OS | -0.05 | 0.00 | 0.00 | 0.19 | 0.00 |
| AL | -0.16 | **1.00** | -0.75 | -0.97 | 0.00 |
| CS | 0.01 | -0.06 | 0.05 | 0.08 | **-1.00** |
| CF | **0.99** | 0.00 | 0.00 | 0.00 | 0.00 |
| Eigenvalue ($\lambda$) | 15.74 | 11.65 | 10.14 | 9.28 | 3.84 |
| Half-life ($t_{1/2} = \frac{\ln 2}{\lambda}$) | 0.04 | 0.06 | 0.07 | 0.07 | 0.18 |
| Tree Height (%) | 4.40 | 5.95 | 6.84 | 7.47 | 18.03 |

When analysing long-term evolutionary tempos using half-lives, we found clear differences among clades. Mammals showed extremely rapid adaptation, with half-lives spanning only 0.66–2.26% of tree height (Table 1a). Reptiles exhibited intermediate tempos, with half-lives of 4.4–18.03% (Table 1c). In contrast, birds evolved much more slowly, with the first four modes ranging from 5.55% to 112%, and the fifth mode—dominated by clutch size—having an effectively infinite half-life, consistent with near-neutral evolution (Table 1b).

Most evolutionary modes (characterized by eigenvectors) in mammals and birds were dominated by a single trait, indicating largely independent axes of long-term adaptation (Table 1). For example, in mammals, all modes but one 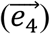 were associated primarily with a single trait (Table 1a). In birds, most modes were similarly trait-specific, except for one mode 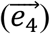 that combined offspring size and development time with opposing loadings (Table 1b). Reptilian modes were more composite: 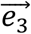 was primarily a combination of adult lifespan and development time, while 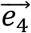 involved adult lifespan and development time, and offspring size (Table 1c).

## Discussion

Life-history evolution is fundamentally complex, driven by interacting genetic, physiological, and ecological mechanisms that shape how organisms allocate limited resources among tasks (De Jong 1993; Stearns 1992). Such a constraint means that traits can rarely evolve independently. Instead, they may coevolve as integrated networks (Del Giudice 2020; Stott *et al*. 2024; Svensson *et al*. 2021; Walsh & Blows 2009). Our study tested this expectation and revealed further complications in life history evolution across amniotes. For example, we showed that the association between clutch size and clutch frequency, previously suggested to be a reflection of direct interaction (Stearns 1992), is actually mediated by offspring size and/or development time (Figures 3 and 4). Similar multivariate associations for other traits have been revealed in marine endotherms, where gestation/incubation time affects maximum longevity through correlated traits such as body size, locomotor morphology and encephalization (Sol *et al*. 2025). Likewise, a three-way trade-off among juvenile mortality, adult mortality and body size was identified in vertebrates (Brooks *et al*. 2025).

We found trait coevolution operated at two distinct timescales: instantaneous vs. long-term (macroevolutionary) couplings. On the short-term scale, we found that the fast– slow continuum identified by the first PCA axis in our analysis only partially captures the multivariate structure of life-history variation (Del Giudice 2020; Stott *et al*. 2024). PCA recovers a broad axis of covariation, but the OU framework shows that this axis conceals important asymmetries: lifespan, development time, and offspring size coevolve tightly, yet clutch size and frequency, despite loading together on the fast– slow axis, lack direct evolutionary coupling (Figure 4). This pattern likely arises because the former three traits are genetically and developmentally related. Larger offspring require longer maturation and are typically linked to delayed senescence and extended survival (Crews & Bogin 2010). In contrast, clutch size and frequency are governed by distinct constraints; therefore, they may covary statistically along the fast– slow axis without being causally linked through direct coevolution.

The long-term patterns of coevolution, revealed via the eigen-decomposition of the ***A*** matrix, further confirm the independence between clutch size and frequency. Clutch frequency dominates the fastest evolutionary mode, while clutch size dominates the slowest, indicating that the rate of reproductive events and the magnitude of investment per event evolve along distinct and temporally decoupled trajectories (Table 1). This divergence likely reflects contrasting constraints: clutch frequency is highly responsive to ecological conditions such as seasonality and resource availability and therefore evolves rapidly (Boutin 1990; Husby *et al*. 2009), whereas clutch size is more tightly constrained by anatomical (e.g., pelvic morphology), developmental and physiological limits on offspring production (e.g., offspring viability, yolk provisioning), thus evolves slowly (Jetz *et al*. 2008; Sinervo & Licht 1991).

Moreover, our findings on instantaneous versus long-term coevolution highlight that variation in life-history strategies can arise from processes operating across multiple temporal scales. We thus suggest a useful complementary approach to understanding how evolutionary constraints act differently across these scales is to expand the comparison to intraspecific variations (Agrawal 2020; Chang *et al*. 2024; Moiron *et al*. 2020; Van De Walle *et al*. 2023). Such cross-scale comparisons can reveal how short-term ecological and demographic processes at intraspecific scales translate into long-term evolutionary trajectories. Recent efforts to compile individual-level life-history databases (Haave-Audet *et al*. 2022; Van De Walle *et al*. 2023) represent a critical step toward bridging micro- and macroevolutionary perspectives in understanding life history evolution (Rolland *et al*. 2023).

We also found clade-specific interactions among traits, reflecting how life strategies are shaped by clade-specific physiological and ecological constraints. For instance, in birds, the positive association between adult lifespan (AL) and clutch size (CS) likely reflects determinate growth: once somatic growth ceases, adults can allocate resources to both survival and reproduction (Sæther & Bakke 2000). Longer-lived birds also gain breeding experience, improve territory defense, and forage more efficiently, enabling larger clutches without compromising survival (Healy *et al*. 2014; Sæther 1988). In mammals, however, gestation and lactation impose strict physiological constraints on litter size, largely decoupling reproduction from lifespan (Charnov & Ernest 2006; Speakman 2008). By contrast, reptiles with indeterminate growth face a persistent trade-off between growth, survival, and reproduction: species with large clutches often experience higher mortality, favoring a “live fast, die young” strategy, whereas long-lived species spread reproductive effort across years, producing fewer but more sustainable clutches (Shine 2005; Shine & Charnov 1992; Warner & Shine 2008). These differences are unlikely to result from sampling bias, as several recovered relationships align with established life-history theory.

Looking forward, there are several avenues that could further advance our understanding of how life-history traits coevolve across diverse clades. For example, expanding taxonomic and intraspecific sampling—particularly for underrepresented avian lineages—would improve the precision of parameter estimates and strengthen tests of cross-clade patterns. In parallel, methodological developments that extend the multivariate Ornstein–Uhlenbeck (OU) framework to incorporate multiple selective optima or context-dependent regimes would provide a more realistic depiction of the adaptive landscapes underlying life-history evolution (Cooper *et al*. 2016). Together, these improvements would move comparative life history research toward a more integrative, multi-scale framework linking individual-level processes to macroevolutionary diversification.

## Acknowledgements

We are grateful to all data collectors and contributors of the Amniote and VertLife databases. We also thank Prof. Shucun Sun and Prof. Xinqiang Xi of Nanjing University, as well as Prof. Shilu Zheng of Xiamen University, for their valuable comments and insightful discussions on the manuscript. This work was supported by the *Yuxiu Young Scholars Program* of Nanjing University, awarded to Zhang Yang.

