## Supplementary figures for "Multivariate Coevolution Shapes Life-History Strategies Across Amniotes"

#### Supplementary materials

#### Section 1: Extended figures used to support findings in the main text.

a

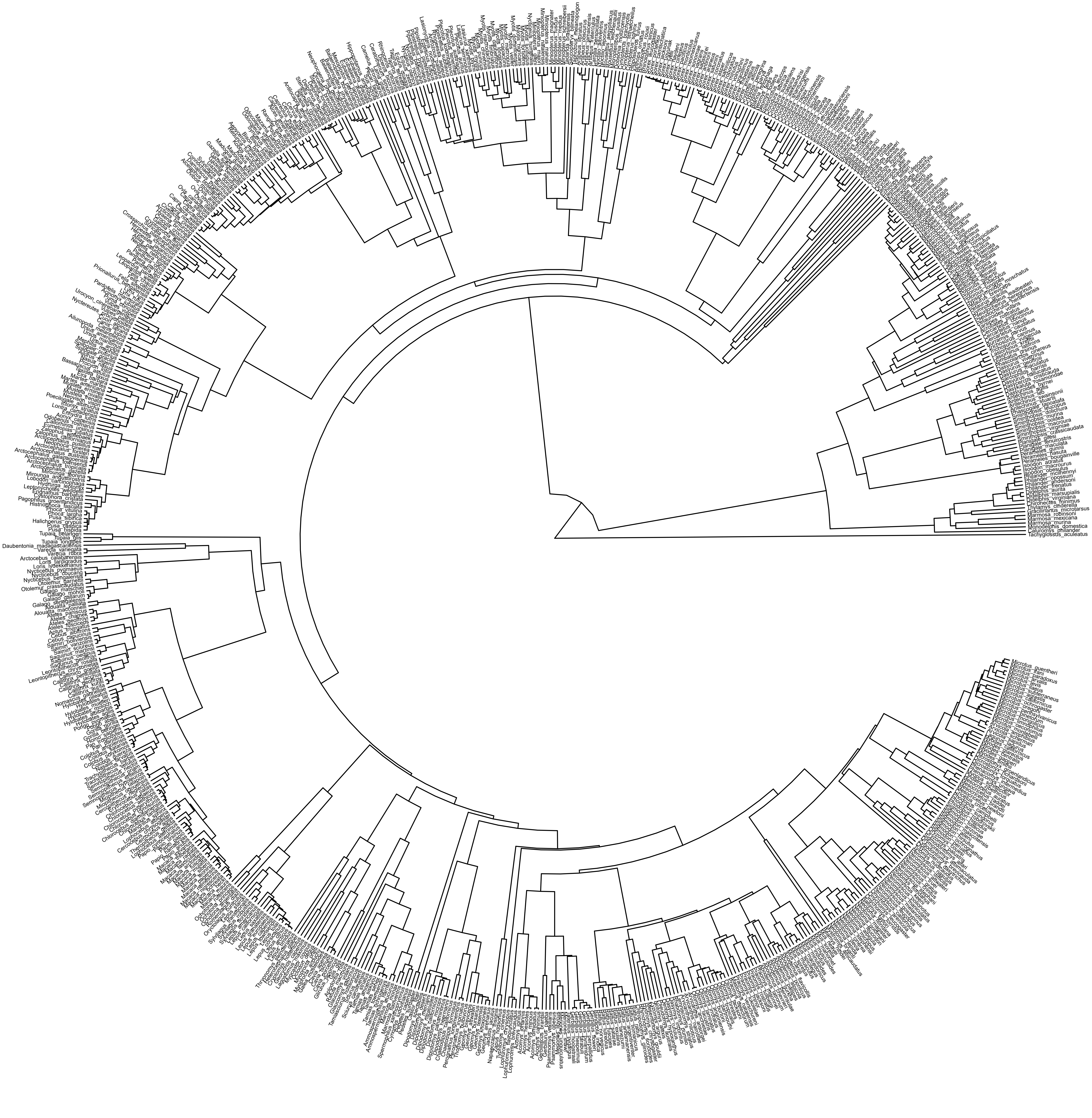

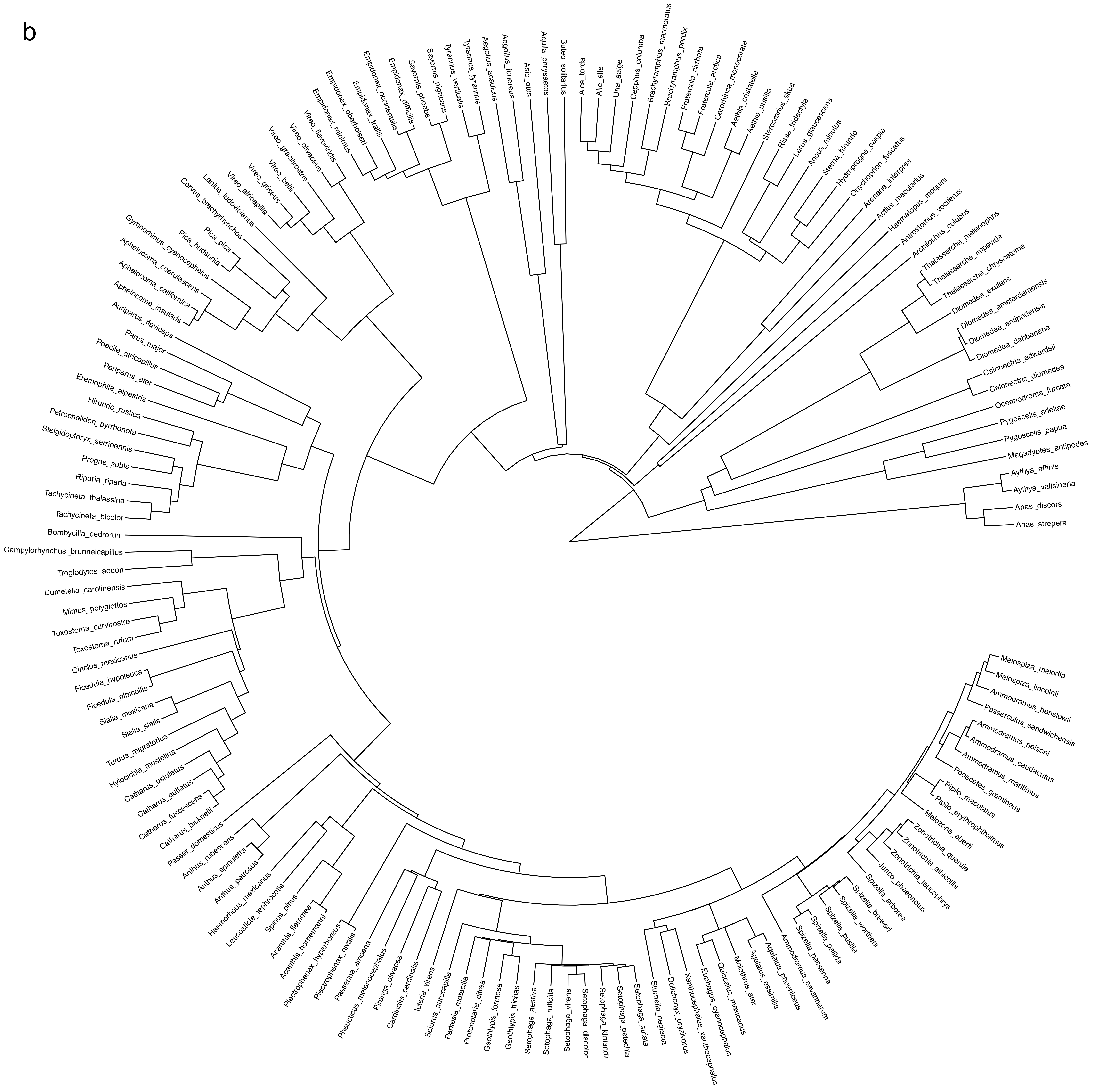
b
Auriparus_flaviceps
Aphelocoma_insularis
Aphelocoma_californica
Aphelocoma_coerulescens
Gymnorhinus_cyanocephalus
Pica_hudsonia
Pica_pica
Corvus_brachyrhynchos
Lanius_ludovicianus
Vireo_atricapilla
Vireo_griseus
Vireo_bellii
Vireo_gracilirostris
Vireo_olivaceus
Vireo_flavoviridis
Empidonax_minimus
Empidonax_oberholseri
Empidonax_traillii
Empidonax_occidentalis
Empidonax_difficilis
Sayornis_phoebe
Sayornis_nigricans
Tyrannus_verticalis
Tyrannus_tyrannus
Aegolius_acadicus
Aegolius_funereus
Asio_otus
Aquila_chrysaetos
Buteo_solitarius
Alca_torda
Alle_alle
Uria_aalge
Cepphus_columba
Brachyramphus_marmoratus
Brachyramphus_perdix
Fratercula_cirrhata
Fratercula_arctica
Cerorhinca_monocerata
Aethia_cristatella
Aethia_pusilla
Stercorarius_skua
Rissa_tridactyla
Larus_glaucescens
Anous_minutus
Sterna_hirundo
Hydroprogne_caspia
Onychoprion_fuscatus
Arenaria_interpres
Actitis_macularius
Haematopus_moquini
Antrostomus_vociferus
Archilochus_colubris
Thalassarche_melanophris
Thalassarche_impavida
Thalassarche_chrysostoma
Diomedea_exulans
Diomedea_amsterdamensis
Diomedea_antipodensis
Diomedea_dabbenena
Calonectris_edwardsii
Calonectris_diomedea
Oceanodroma_furcata
Pygoscelis_adeliae
Pygoscelis_papua
Megadyptes_antipodes
Aythya_affinis
Aythya_valisineria
Anas_discors
Anas_strepera
Melospiza_melodia
Melospiza_lincolnii
Ammodramus_henslowii
Passerculus_sandwichensis
Ammodramus_nelsoni
Ammodramus_caudacutus
Ammodramus_maritimus
Pooecetes_gramineus
Pipilo_maculatus
Pipilo_erythrophthalmus
Melozone_aberti
Zonotrichia_querula
Zonotrichia_albicollis
Zonotrichia_leucophrys
Junco_phaeonotus
Spizella_arborea
Spizella_breweri
Spizella_wortheni
Spizella_pusilla
Spizella_pallida
Spizella_passerina
Ammodramus_savannarum
Agelaius_phoeniceus
Agelaius_assimilis
Molothrus_ater
Quiscalus_mexicanus
Euphagus_cyanocephalus
Xanthocephalus_xanthocephalus
Dolichonyx_oryzivorus
Sturnella_neglecta
Setophaga_striata
Setophaga_petechia
Setophaga_kirtlandii
Setophaga_discolor
Setophaga_virens
Setophaga_ruticilla
Setophaga_aestiva
Geothlypis_trichas
Geothlypis_formosa
Protonotaria_citrea
Parkesia_motacilla
Seiurus_aurocapilla
Icteria_virens
Cardinalis_cardinalis
Piranga_olivacea
Pheucticus_melanocephalus
Passerina_amoena
Plectrophenax_nivalis
Plectrophenax_hyperboreus
Acanthis_hornemanni
Acanthis_flammea
Spinus_pinus
Leucosticte_tephrocotis
Haemorhous_mexicanus
Anthus_petrosus
Anthus_spinoletta
Anthus_rubescens
Passer_domesticus
Catharus_bicknelli
Catharus_fuscescens
Catharus_guttatus
Catharus_ustulatus
Hylocichla_mustelina
Turdus_migratorius
Sialia_sialis
Sialia_mexicana
Ficedula_albicollis
Ficedula_hypoleuca
Cinclus_mexicanus
Toxostoma_rufum
Toxostoma_curvirostre
Mimus_polyglottos
Dumetella_carolinensis
Troglodytes_aedon
Campylorhynchus_brunneicapillus
Bombycilla_cedrorum
Tachycineta_bicolor
Tachycineta_thalassina
Riparia_riparia
Progne_subis
Stelgidopteryx_serripennis
Petrochelidon_pyrrhonota
Hirundo_rustica
Eremophila_alpestris
Periparus_ater
Poecile_atricapillus
Parus_major

c

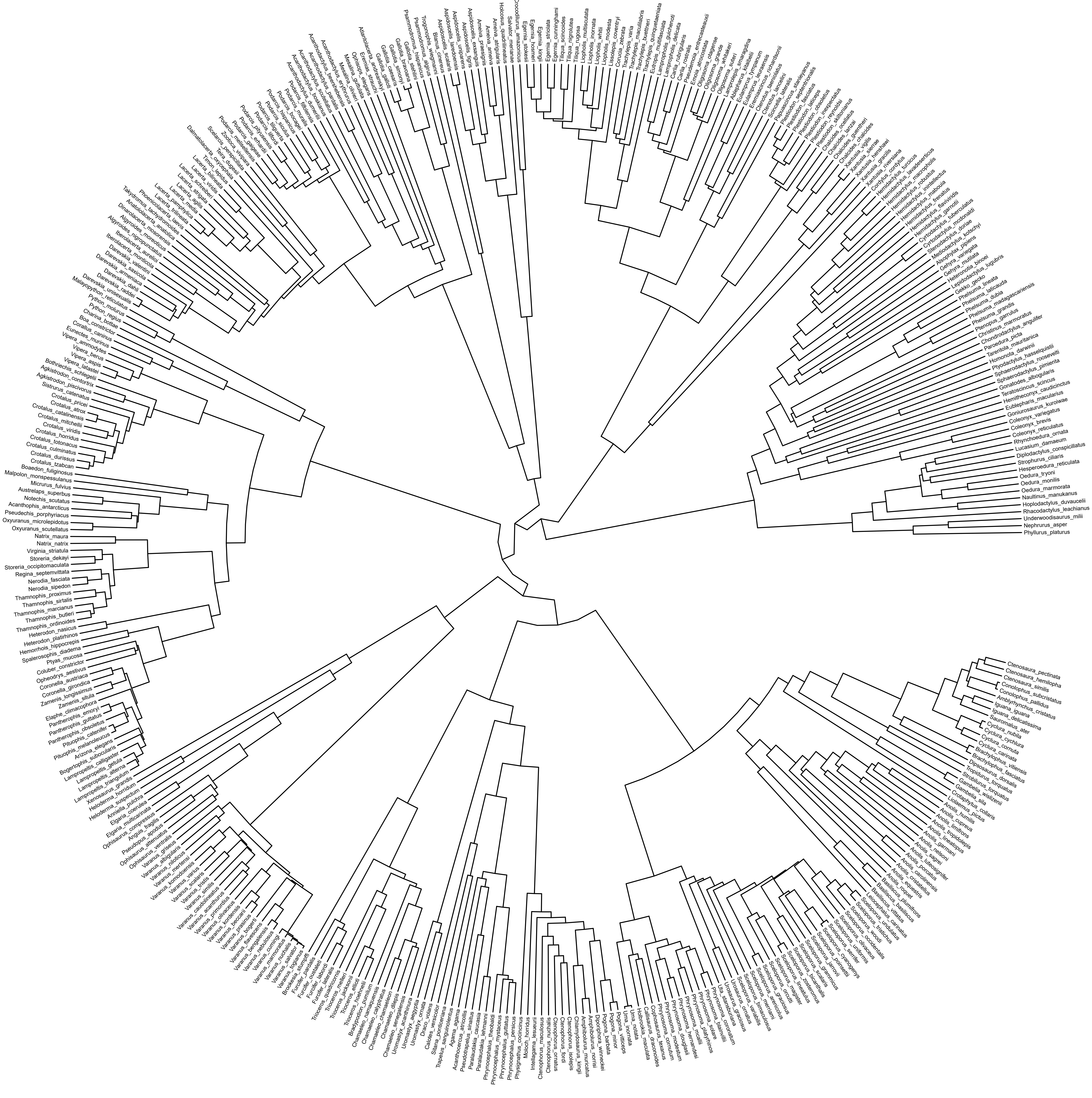

**Figure S1.** Phylogenetic trees of the species represented in our dataset, including (a) 774 mammalian species, (b) 156 avian species, and (c) 411 reptilian species. For each taxon, we pruned published phylogenetic trees from the Vertlife database (https://vertlife.org/; accessed June 25, 2025) to reflect the shared evolutionary histories of the species included in our analyses.

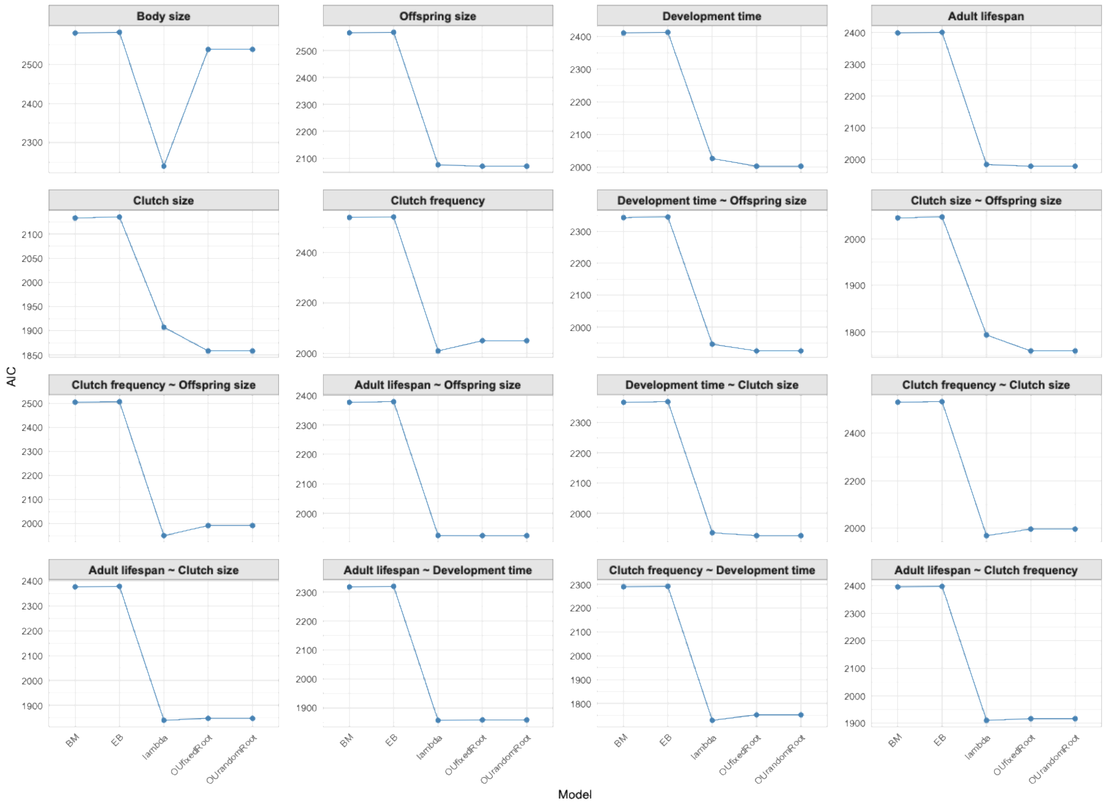


**Figure S2.** AIC comparison for different models and life-history traits across mammals. Each plot illustrates the AIC values for various models in relation to individual life-history trait and their pairwise relationships. The models assessed include BM, EB, lambda, OUfixedRoot, and OUrandomRoot, emphasizing the variation in model fit across species.

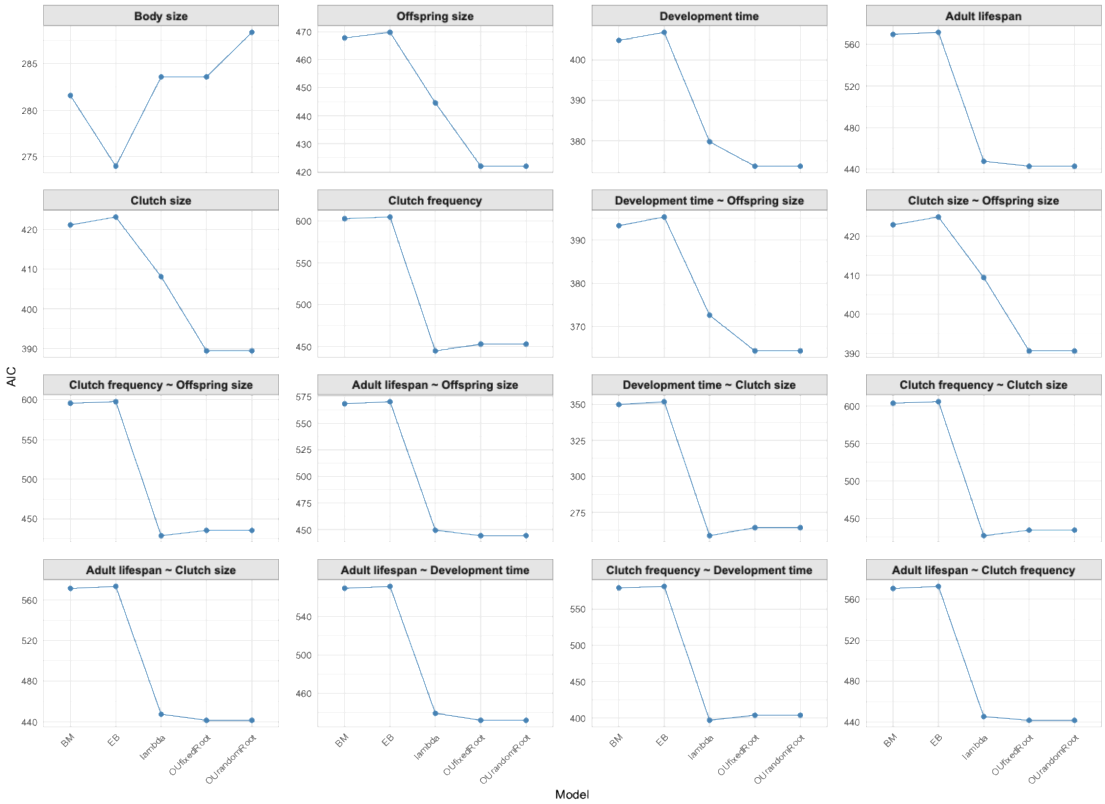


**Figure S3.** AIC comparison for different models and life-history traits across birds. Each plot illustrates the AIC values for various models in relation to individual life-history trait and their pairwise relationships. The models assessed include BM, EB, lambda, OUfixedRoot, and OUrandomRoot, emphasizing the variation in model fit across species.

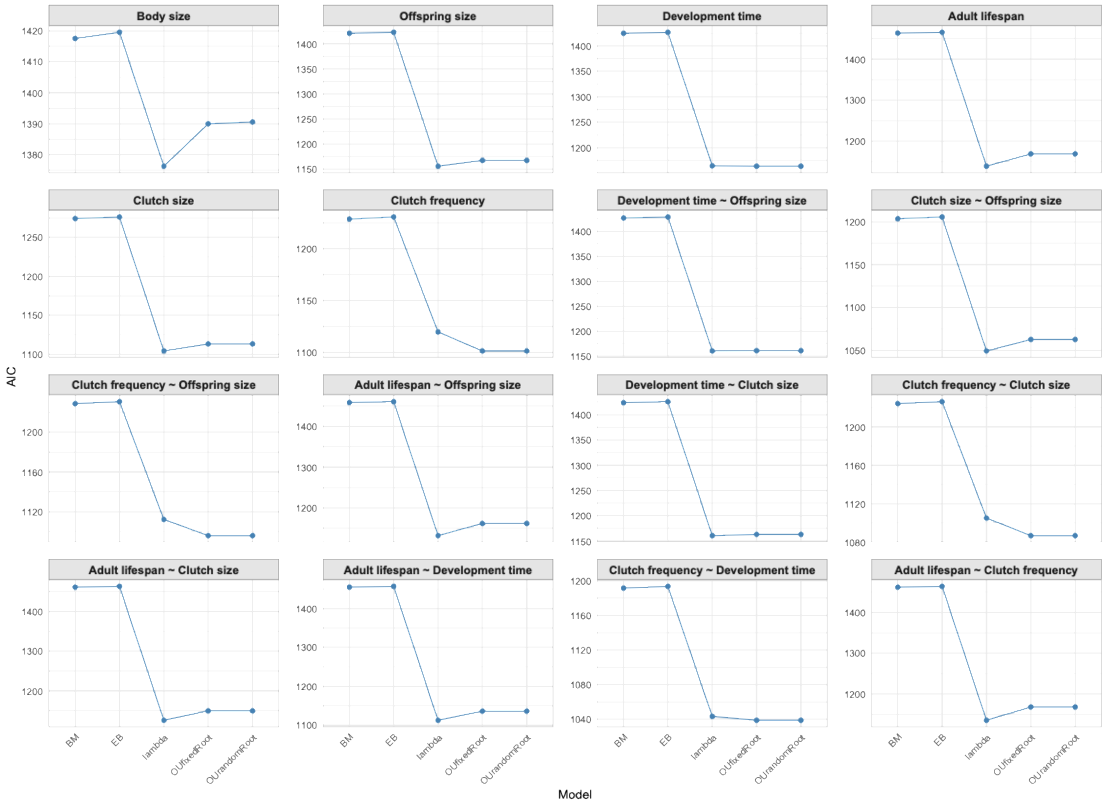


**Figure S4.** AIC comparison for different models and life-history traits across reptiles. Each plot illustrates the AIC values for various models in relation to individual life-history trait and their pairwise relationships. The models assessed include BM, EB, lambda, OUfixedRoot, and OUrandomRoot, emphasizing the variation in model fit across species.

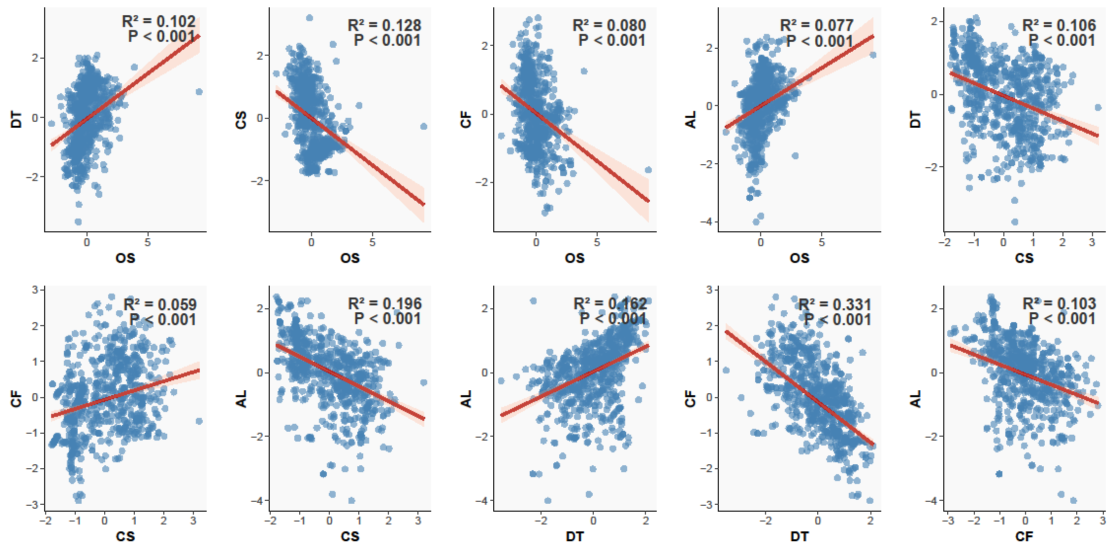


**Figure S5.** Pairwise correlations among life-history traits across 774 mammalian species based on PGLS models. Each scatterplot shows the relationship between two traits, with individual species represented by blue dots. The red line indicates the best-fit regression, with shaded areas representing confidence intervals. Reported statistics include the coefficient of determination ($R^2$) and significance (*P*) for each relationship. Trait abbreviations: OS = offspring size; DT = development time; AL = adult lifespan; CS = clutch size; CF = clutch frequency.

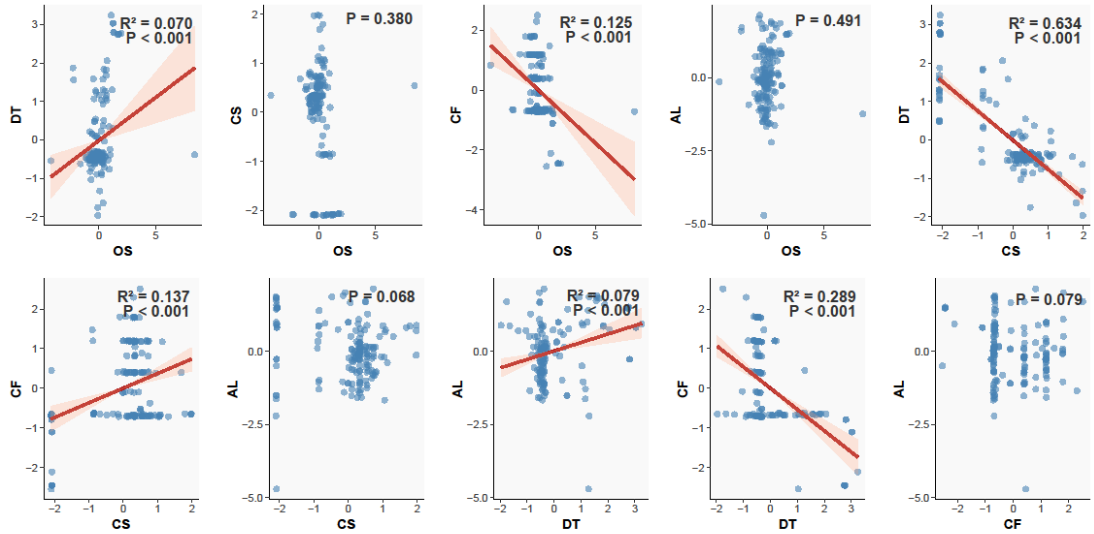


**Figure S6.** Pairwise correlations among life-history traits across 156 avian species based on PGLS models. Each scatterplot shows the relationship between two traits, with individual species represented by blue dots. The red line indicates the best-fit regression, with shaded areas representing confidence intervals. Reported statistics include the coefficient of determination ($R^2$) and significance (*P*) for each relationship. Trait abbreviations: OS = offspring size; DT = development time; AL = adult lifespan; CS = clutch size; CF = clutch frequency.

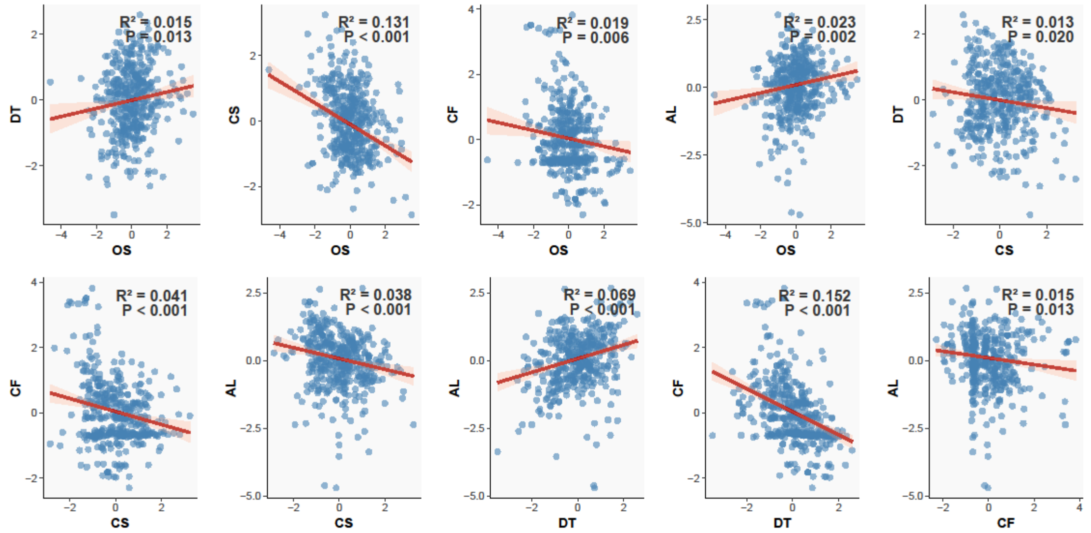


**Figure S7.** Pairwise correlations among life-history traits across 411 reptilian species based on PGLS models. Each scatterplot shows the relationship between two traits, with individual species represented by blue dots. The red line indicates the best-fit regression, with shaded areas representing confidence intervals. Reported statistics include the coefficient of determination ($R^2$) and significance (*P*) for each relationship. Trait abbreviations: OS = offspring size; DT = development time; AL = adult lifespan; CS = clutch size; CF = clutch frequency.

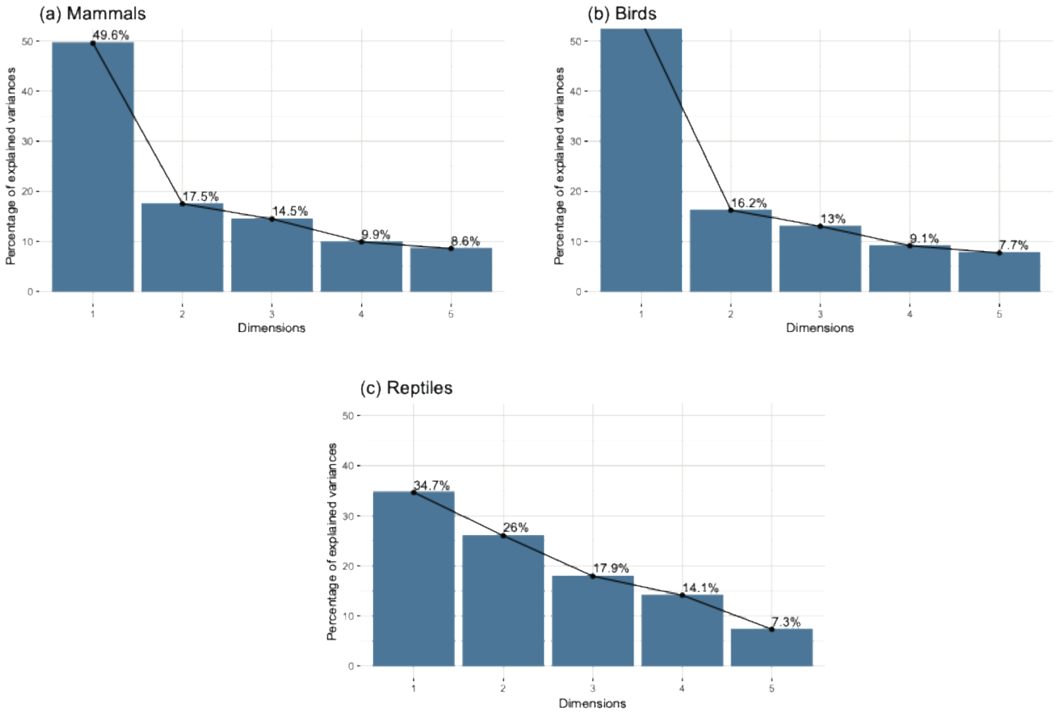


**Figure S8.** Scree plots from phylogenetically informed PCA (pPCA) for (a) mammals, (b) birds, and (c) reptiles. The plots display the percentage of variance explained by the first five principal components, indicating the relative contribution of each axis to overall variation in life-history traits.

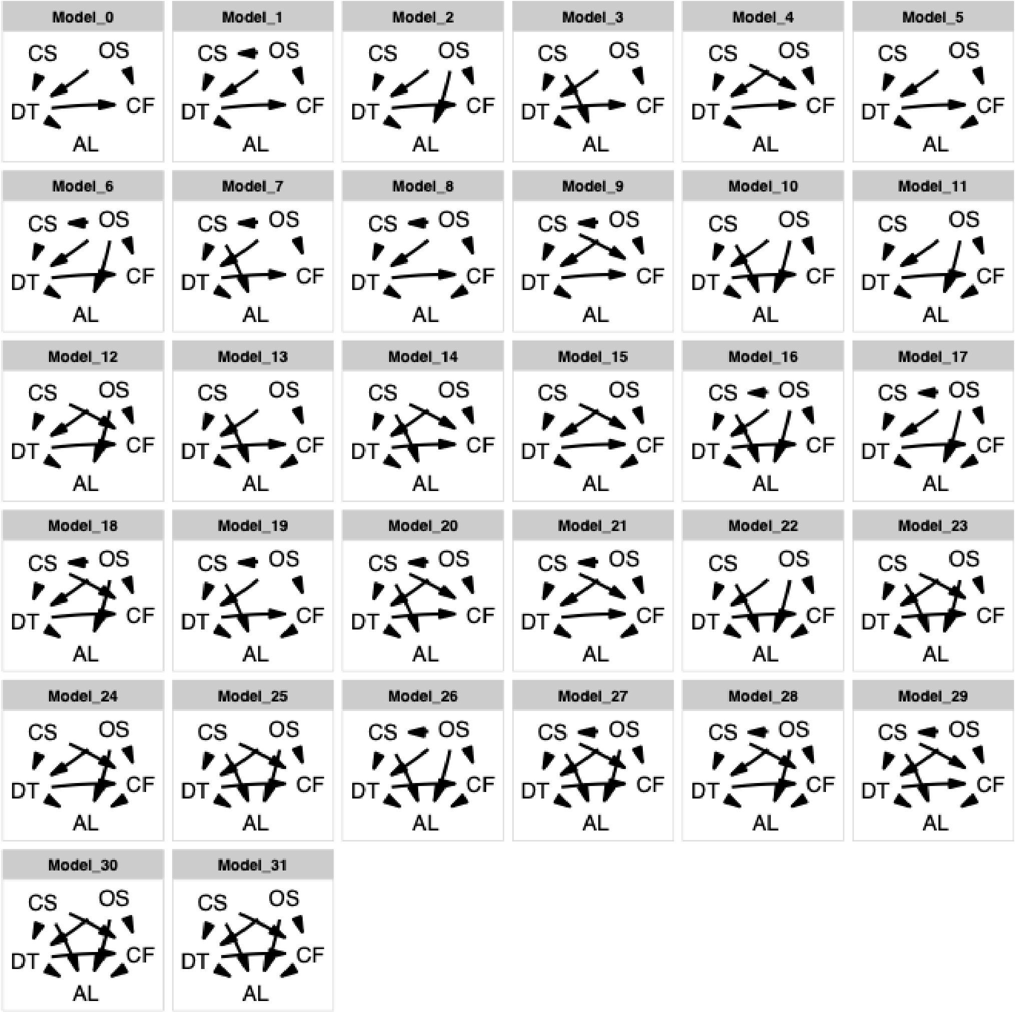


**Figure S9.** Directed Acyclic Graph (DAG) models showing the direct relationships between life-history traits across species. Each panel represents a different model (Model 0 to Model 31), illustrating how traits such as clutch size (CS), offspring size (OS), development time (DT), adult lifespan (AL), and clutch frequency (CF) are interconnected. The arrows simply indicate the pairwise relationships between the variables, without implying causality.

### Section 2: Robustness tests on the findings in the main text.

#### Part 1: Robustness tests on pPCA Patterns

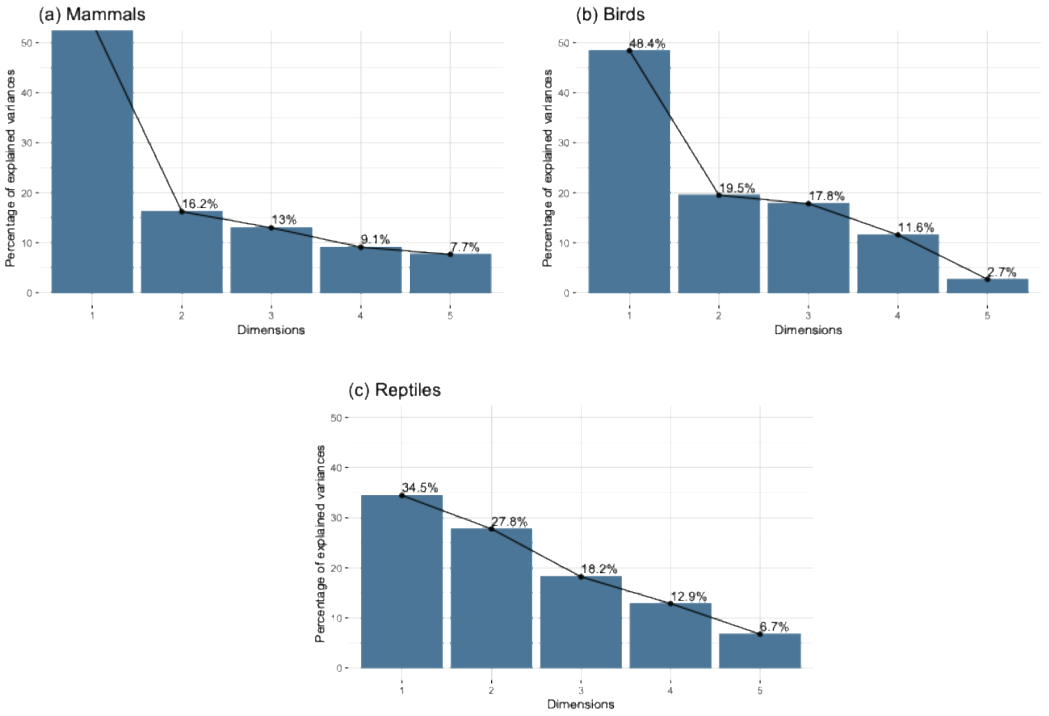


**Figure S10.** Scree plots from PCA for (a) mammals, (b) birds, and (c) reptiles. The plots display the percentage of variance explained by the first five principal components, indicating the relative contribution of each axis to overall variation in life-history traits.

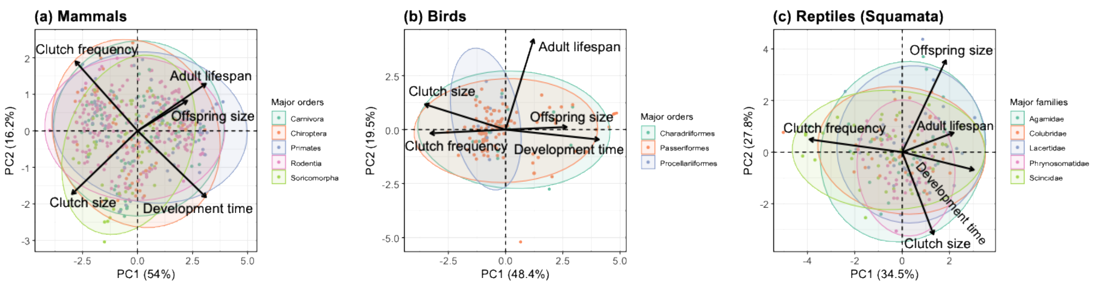


**Figure S11.** Biplots of PCA illustrating the covariations among five life-history traits in (a) 774 mammalian, (b) 156 avian, and (c) 411 reptilian (squamate) species. Arrows represent the loadings of five key traits on the first two principal components (PCs). Across all groups, PC1 primarily captures the fast-slow continuum of life-history strategies, while PC2 reflects secondary dimensions of trait covariation. Each point corresponds to a species, with colors denoting major clades, including five mammalian orders, three avian orders, and five squamate families. Ellipses summarize the distribution of each clade in multivariate trait space, emphasizing both shared patterns and clade-specific divergences. Collectively, the first two axes explain 70.2% of the total variation in mammals, 67.9% in birds, and 62.3% in reptiles. Our analysis confirmed that the patterns identified in pPCA were also supported by PCA, reinforcing the reliability of the life-history trait structures across different taxa. This consistency underscores the universal nature of the trade-offs driving life-history variation, irrespective of species' taxonomic classification.

#### Part 2: Robustness tests on the patterns of PPA

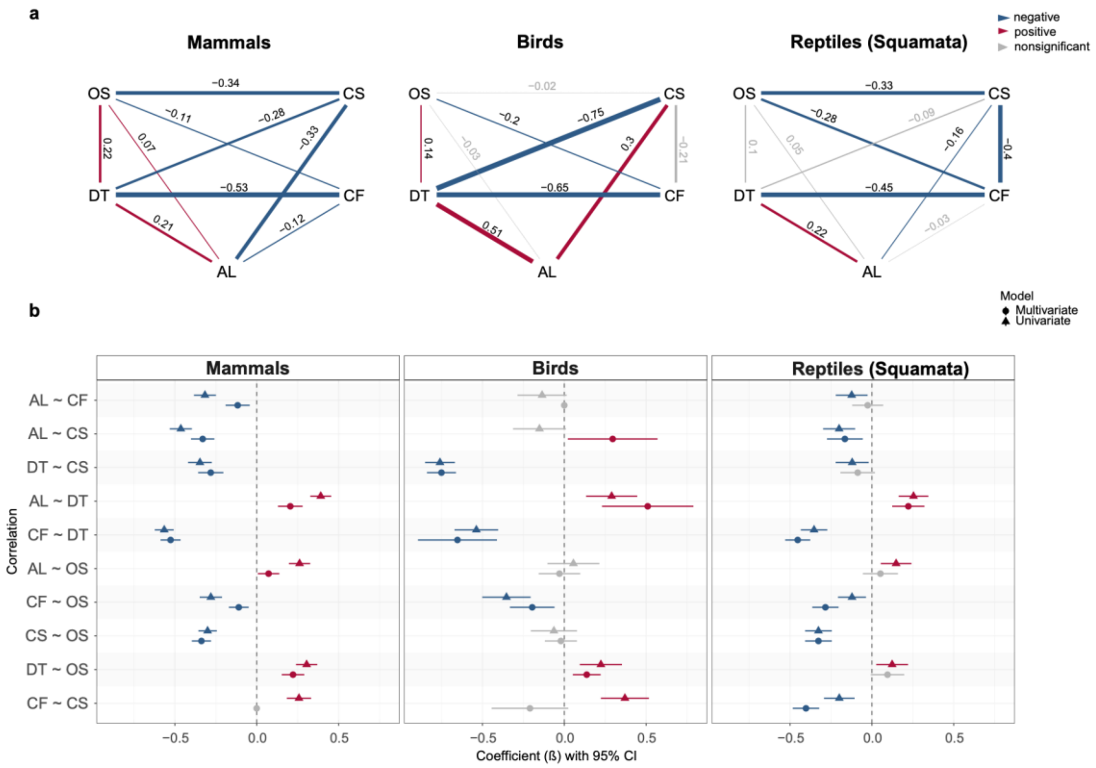


**Figure S12.** (a) Averaged best models of phylogenetic path analysis (PPA) under Pagel's λ model for mammals, birds, and reptiles. Red and blue lines represent significant positive and negative associations between pairs of traits, respectively, with the thickness of each line reflecting the magnitude of their standardized path coefficients. Gray lines indicate non-significant relationships between traits. The values adjacent to the lines denote the standardized average path coefficients. (b) Comparison of standardized regression coefficients from univariate (PGLS) and multivariate (PPA) models. Forest plots show regression coefficients with ± 95% confidence intervals for each pair of traits. Red and blue circles or triangles represent significant positive and negative associations, respectively, while gray circles or triangles indicate non-significant relationships. Triangular markers represent the coefficients derived from the PPA, whereas circular markers correspond to those from the PGLS models. Trait abbreviations: OS = offspring size; DT = development time; AL = adult lifespan; CS = clutch size; CF = clutch frequency. Overall, the results were highly consistent with those obtained under the OU model, showing only minor differences across species groups. It is suggested that the observed life-history patterns are robust to model specification, with only limited variation in the strength or direction of certain trait relationships.

#### Part 3: Robustness tests on the patterns of mvOU model in birds

(a)

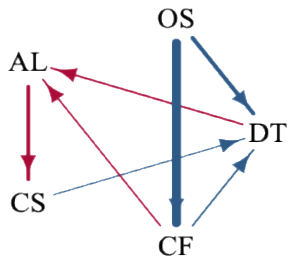


(b)

**Bird (imputated)**

| Trait / Metric | $\vec{e1}$ | $\vec{e2}$ | $\vec{e3}$ | $\vec{e4}$ | $\vec{e5}$ |
|---|---|---|---|---|---|
| DT | -0.06 | -0.04 | 0.00 | 0.00 | **-1.00** |
| OS | 0.48 | 0.00 | 0.00 | 0.00 | 0.00 |
| AL | -0.05 | 0.12 | 0.65 | 0.00 | 0.08 |
| CS | -0.01 | 0.06 | 0.76 | **-1.00** | -0.03 |
| CF | -0.87 | **0.99** | 0.00 | 0.00 | 0.00 |
| Eigenvalue ($\lambda$) | 8.00 | 5.81 | 4.39 | 3.31 | 0.19 |
| Half-life ($t_{1/2} = \frac{ln2}{\lambda}$) | 0.09 | 0.12 | 0.16 | 0.21 | 3.71 |
| Tree Height (%) | 8.66 | 11.94 | 15.77 | 20.97 | 370.64 |

(c)

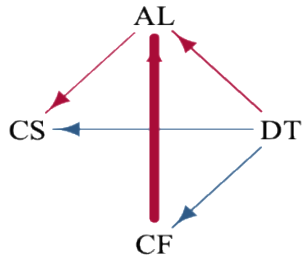


(d)

**Bird (without offspring size)**

| Trait / Metric | $\vec{e1}$ | $\vec{e2}$ | $\vec{e3}$ | $\vec{e4}$ |
|---|---|---|---|---|
| DT | 0.00 | 0.01 | 0.00 | **1.00** |
| AL | 0.00 | 0.73 | 0.00 | 0.00 |
| CS | 0.00 | 0.00 | **1.00** | 0.00 |
| CF | **1.00** | -0.68 | 0.00 | 0.00 |
| Eigenvalue ($\lambda$) | 66.43 | 62.82 | 20.09 | 5.65 |
| Half-life ($t_{1/2} = \frac{ln2}{\lambda}$) | 0.01 | 0.01 | 0.03 | 0.12 |
| Tree Height (%) | 1.04 | 1.10 | 3.45 | 12.28 |

**Figure S13.** (a, c) Directed causal networks showing the inferred structure of coevolutionary interactions among life-history traits for birds, based on (a) the phylogenetically imputed dataset and (c) the dataset excluding offspring size. Arrows represent coevolutionary links where a change in trait $j$ drives evolutionary change in trait $i$. Red arrows indicate pull towards an optimal value, while blue arrows indicate push away from an optimum. Coefficients on arrows represent the strength of the coevolutionary coupling. (b, d) Eigen-decomposition of the $\boldsymbol{A}$ matrix for each avian dataset is shown, with panel b corresponding to the imputed dataset and panel d to the complete-case dataset. Each eigenvector represents an independent axis of adaptation in the multi-trait space, with its corresponding eigenvalue ($\lambda$) determining the rate of adaptation toward the optimum. The rate of adaptation is then summarized by the evolutionary half-life ($ln(2)/\lambda$) and its proportion of the total tree height. Trait loadings for each eigenvector are displayed. For a given eigenvector, the trait that dominates the adaptive mode is highlighted in bold. Trait abbreviations: OS = offspring size; DT = development time; AL = adult lifespan; CS = clutch size; CF = clutch frequency. The causal architectures recovered here reinforce the main-text findings that clutch size and clutch frequency evolve independently at both instantaneous and long-term timescales, with clutch frequency adapting rapidly while clutch size evolves slowly over macroevolutionary time.

#### Section 3: PGLS, pPCA and PPA analyses without body size control

We performed phylogenetic generalized least squares (PGLS), phylogenetically informed principal component analysis (pPCA), and phylogenetic path analysis (PPA) without controlling for body size (BS), treating it as a separate predictor in the models. In this framework, the remaining life-history traits were analyzed independently of BS to assess their interrelationships. Our results confirmed that BS had significant associations with all five other life-history traits across the studied taxa, underscoring its broad impact on life-history variation.

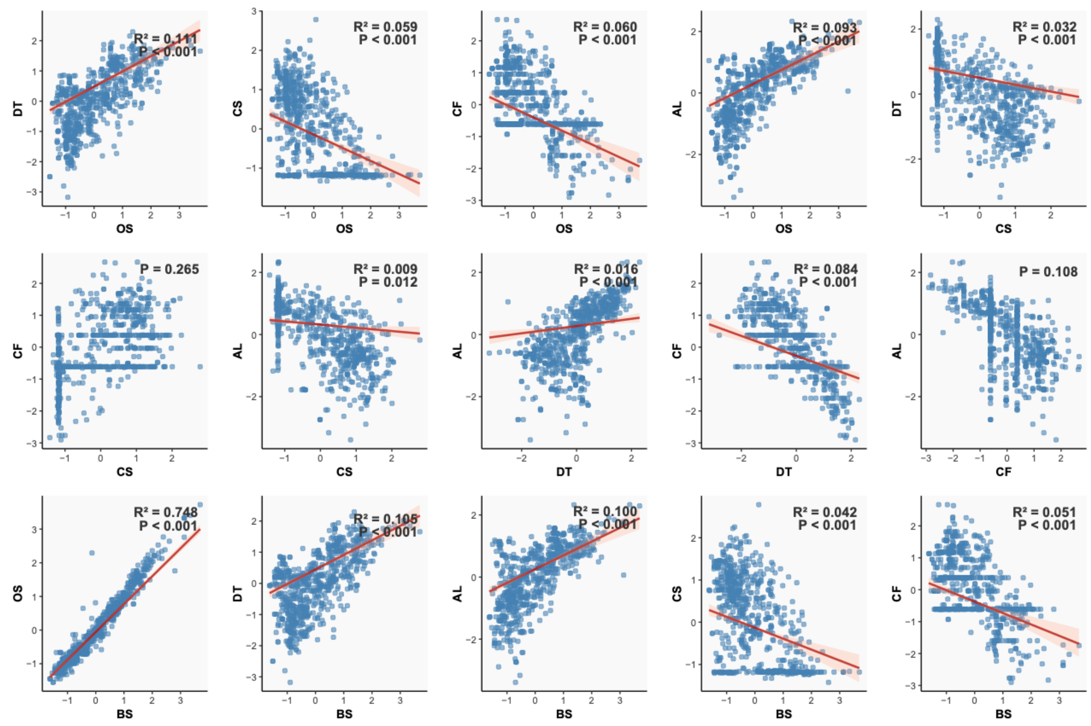


**Figure S14.** Pairwise correlations among life-history traits across 774 mammalian species based on PGLS models. Each scatterplot shows the relationship between two traits, with individual species represented by blue dots. The red line indicates the best-fit regression, with shaded areas representing confidence intervals. Reported statistics include the coefficient of determination ($R^2$) and significance (P) for each relationship. Trait abbreviations: BS = Body size; OS = offspring size; DT = development time; AL = adult lifespan; CS = clutch size; CF = clutch frequency.

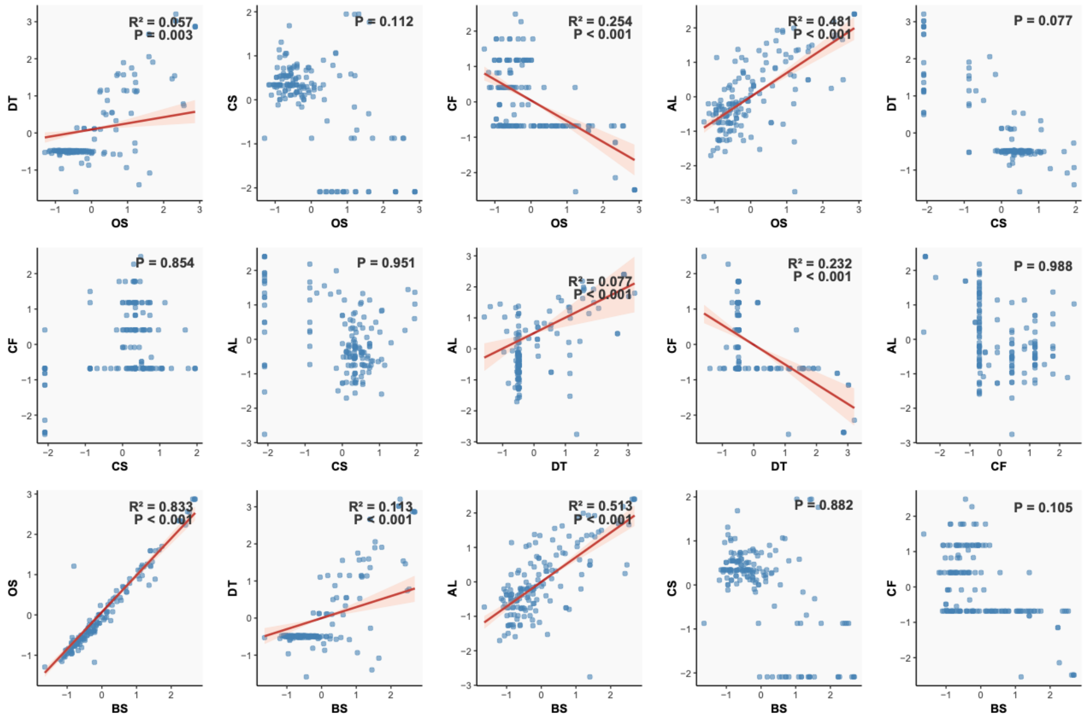


**Figure S15.** Pairwise correlations among life-history traits across 156 avian species based on PGLS models. Each scatterplot shows the relationship between two traits, with individual species represented by blue dots. The red line indicates the best-fit regression, with shaded areas representing confidence intervals. Reported statistics include the coefficient of determination ($R^2$) and significance (*P*) for each relationship. Trait abbreviations: BS = Body size; OS = offspring size; DT = development time; AL = adult lifespan; CS = clutch size; CF = clutch frequency.

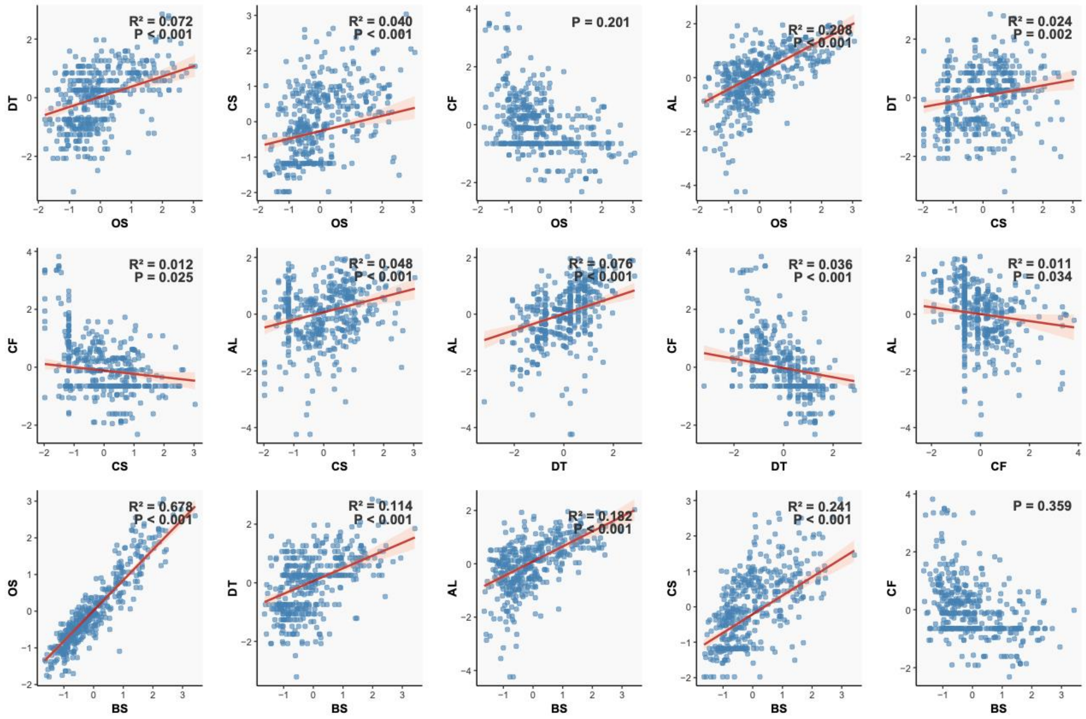


**Figure S16.** Pairwise correlations among life-history traits across 411 reptilian species based on PGLS models. Each scatterplot shows the relationship between two traits, with individual species represented by blue dots. The red line indicates the best-fit regression, with shaded areas representing confidence intervals. Reported statistics include the coefficient of determination ($R^2$) and significance (*P*) for each relationship. Trait abbreviations: BS = Body size; OS = offspring size; DT = development time; AL = adult lifespan; CS = clutch size; CF = clutch frequency.

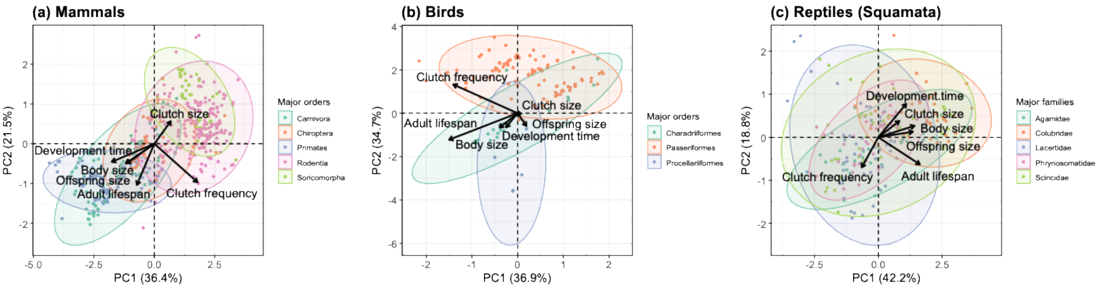


**Figure S17.** Biplots of phylogenetically informed PCA illustrating the covariations among body size and five life-history traits in (a) 774 mammalian, (b) 156 avian, and (c) 411 reptilian (squamate) species. Arrows represent the loadings of five key traits on the first two principal components (PCs). Across all groups, PC1 primarily captures the fast-slow continuum of life-history strategies, while PC2 reflects secondary dimensions of trait covariation. Each point corresponds to a species, with colors denoting major clades, including five mammalian orders, three avian orders, and five squamate families. Ellipses summarize the distribution of each clade in multivariate trait space, emphasizing both shared patterns and clade-specific divergences. Collectively, the first two axes explain 57.9% of the total variation in mammals, 71.6% in birds, and 61% in reptiles.

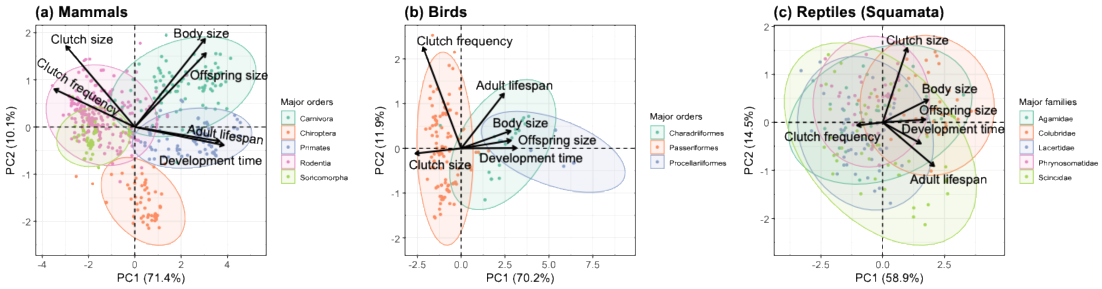


**Figure S18.** Biplots of PCA illustrating the covariations among body size and five life-history traits in (a) 774 mammalian, (b) 156 avian, and (c) 411 reptilian (squamate) species. Arrows represent the loadings of five key traits on the first two principal components (PCs). Across all groups, PC1 primarily captures the fast-slow continuum of life-history strategies, while PC2 reflects secondary dimensions of trait covariation. Each point corresponds to a species, with colors denoting major clades, including five mammalian orders, three avian orders, and five squamate families. Ellipses summarize the distribution of each clade in multivariate trait space, emphasizing both shared patterns and clade-specific divergences. Collectively, the first two axes explain 81.5% of the total variation in mammals, 82.1% in birds, and 73.4% in reptiles.

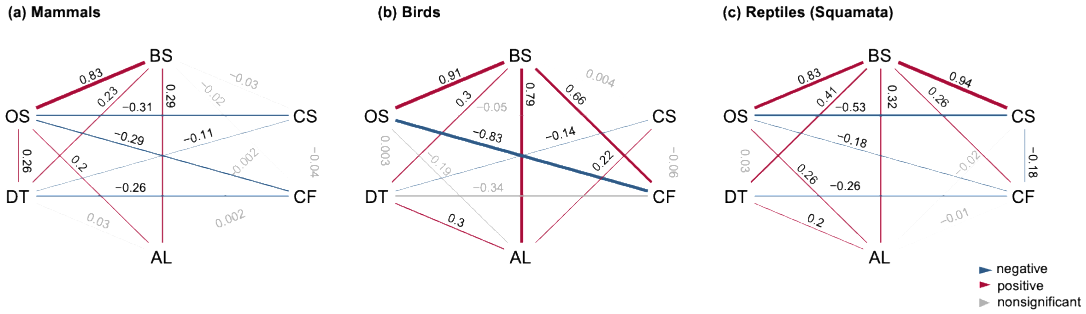


**Figure S19.** Averaged best models of phylogenetic path analysis (PPA) for mammals, birds, and reptiles. Red and blue lines represent significant positive and negative associations between pairs of traits, respectively, with the thickness of each line reflecting the magnitude of their standardized path coefficients. Gray lines indicate non-significant relationships between traits. The values adjacent to the lines denote the standardized average path coefficients. Trait abbreviations: BS = body size; OS = offspring size; DT = development time; AL = adult lifespan; CS = clutch size; CF = clutch frequency.

#### Section 4: Supplementary tables

**Table S1.** Species across three taxa and their life-history traits included in this study.

**Table S2.** Results from phylogenetic generalized least squares (PGLS) analyses across three taxa, illustrating the relationships between pairs of life-history traits. For each trait pair, estimated coefficients, standard errors (SE), *t* values, bootstrap confidence intervals, *p* values, and adjusted $R^2$ values are provided, quantifying the strength and statistical significance of the associations.

**Table S3.** Model comparison of univariate and multivariate PGLS models for life-history trait associations. For each trait treated as the response variable, a full model including the remaining four traits as predictors was fitted, followed by all possible reduced models generated by removing one or more predictors. Model fit was assessed using Akaike Information Criterion (AIC), with the lowest AIC indicating the best-fitting model.

**Table S4.** Models tested in our PPA analyses. In our analyses we included a total of 32 models. Each model in the list represents a different set of previously proposed relationships between the traits, where the arrow (~) indicates a relationship or a path between two life-history traits. Consistent pairwise associations identified across all taxa in the PGLS analyses were incorporated as fixed components in every candidate model. For associations that varied across taxa, we systematically adjusted their inclusion across different models. Trait abbreviations: OS = offspring size; DT = development time; AL = adult lifespan; CS = clutch size; CF = clutch frequency.

**Table S5.** Complete results of the phylogenetic path analysis for mammalian, avian, and reptilian species, based on different evolutionary models. The best models are highlighted in bold, and their averages are summarized in the main text. Here, *k* represents the number of conditional independencies tested, *q* indicates the number of estimated parameters, and C refers to Fisher's C statistic.

**Table S6.** Phylogenetic path analysis (PPA) results for mammals, birds, and reptiles under the Ornstein–Uhlenbeck (OU) model, showing estimated coefficients, standard errors (SE), and 95% confidence intervals for pairwise trait associations.

**Table S7.** Phylogenetic path analysis (PPA) results for mammals, birds, and reptiles under the Pagel's λ model, showing estimated coefficients, standard errors (SE), and 95% confidence intervals for pairwise trait associations.

**Table S8.** Model selection results of multivariate Ornstein–Uhlenbeck (OU) analyses for five life-history traits in mammals, birds, and reptiles under two diffusion matrix. For each candidate model, minimum values of AIC, AICc, SIC, BIC, as well as $R^2$, loglikelihood (LogLik), and degrees of freedom (DOF) are reported to assess model fit. Lower AICc values indicate better model performance, accounting for both goodness-of-fit and model complexity.
