## Supplementary methods for "Multivariate Coevolution Shapes Life-History Strategies Across Amniotes"

### Section 1: Definition of life-history traits

We characterized the life history of mammals, birds and reptiles based on six baseline traits: weaning/fledging/hatching mass, age at sexual maturity, weaning/fledging time, maximum longevity, litter/clutch size and litters/clutches per year (see details in Myhrvold *et al.* 2015). Life-history data were primarily sourced from Myhrvold *et al.* (2015), with mammalian records updated using Bakewell *et al.* (2020) and Capellini *et al.* (2015), and reptile records supplemented with Allen *et al.* (2017) and Bakewell *et al.* (2020).

To facilitate cross-taxonomic comparisons among amniotes, we focused on five key life-history traits for which consistent definitions and broad data availability exist across taxa. These traits capture core aspects of development, reproduction and longevity, and are widely used to quantify life-history strategies (Allen *et al.* 2017; Morrow *et al.* 2020) and to predict population growth dynamics (Sæther *et al.* 2013). The selected traits were defined as follows: (1) Offspring size (OS): Body mass at independence—defined as weaning for mammals, fledging for birds, and hatching for reptiles—measured in grams; (2) Development time (DT): Duration from independence to sexual maturity, measured in days. This period corresponds to the interval from weaning to sexual maturity in mammals, fledging to sexual maturity in birds, and hatching to sexual maturity in reptiles; (3) Adult reproductive lifespan (AL): Time from sexual maturity to death, measured in days, reflecting the period during which individuals can reproduce; (4) Clutch size (CS): Number of offspring produced per reproductive event in mammals, or the number of eggs laid by birds and reptiles; (5) Clutch frequency (CF): Number of litters or clutches produced per year. These five traits provide a standardized, taxonomically comparable framework for quantifying life-history strategies across amniotes. The final dataset comprised 1341 amniote species, including 774 mammals, 156 birds, and 411 reptiles (Table S1).

### Section 2: Controlling for body size

In addition to the five life-history traits mentioned above, we also collected data on adult body size, defined as the adult body mass for each species (measured in grams). Given body size is a fundamental allometric factor shaping most life-history traits and metabolic rates (Beccari *et al.* 2024; Hallmann & Griebeler 2018; Jeschke & Kokko 2009), and our analyses confirmed its associations with five life-history traits (see details in Supplementary Materials section 3), we controlled for its effect in our analyses. Specifically, of the five life-history traits analyzed, each was individually regressed with the log-transformed adult body size using phylogenetic generalized least squares (PGLS) regressions (Martins & Hansen 1997). We extracted the body size-corrected residuals from these regressions for each species (Beccari *et al.* 2024; Hallmann & Griebeler 2018; Jeschke & Kokko 2009), which were then centered and scaled as standardized, size-adjusted life-history traits. Our main analyses were based on these size-adjusted life-history traits.

### Section 3: Construction and pruning of phylogenetic trees

For mammals, we utilized the maximum clade credibility tree (MCC tree) from Upham *et al.* (2019) and pruned it with the "*keep.tip*" function from the R package "*ape*" (version 5.8; Paradis *et al.* 2004), retaining only the species present in our dataset. The avian phylogeny used in this study was pruned based on the dated tree of extant bird species published by Jetz *et al.* (2012), which was built using the Hackett *et al.* (2008) backbone. Because reptiles are paraphyletic, our analyses focused exclusively on

Squamata, the most diverse reptilian order, which comprises more than 95% of extant reptile species (Pincheira-Donoso *et al.* 2013). For this group, we adopted the fully sampled squamate phylogeny constructed by Tonini *et al.* (2016). The final pruned phylogenetic trees for each taxon are presented as dendrograms in the supplementary material (Figure S1).

### Section 4: Evaluating multivariate structure in life-history trait evolution

To assess whether life-history traits are better explained by multivariate rather than bivariate relationships, we compared the goodness-of-fit of univariate and multivariate phylogenetic generalized least squares (PGLS) models. For each focal trait treated as the response variable (Y), we fitted a full model including the remaining four life-history traits as predictors (e.g., AL ~ CS + CF + DT + OS, etc.). We then constructed all possible reduced models by removing one or more predictors (e.g., AL ~ CS + CF + DT, AL ~ CS + CF, etc.), thereby generating the complete set of univariate and multivariate predictor combinations. Model performance was evaluated using Akaike Information Criterion (AIC), with the lowest AIC indicating the best-fitting model (Table S3).

### Section 5: Sensitivity analyses for PCA and PPA

To evaluate whether the baseline patterns identified by phylogenetic-informed principal component analysis (pPCA) were robust to a normal PCA, we repeated the analysis without correcting for phylogenetic non-independence (Figure S11). In both the phylogenetically informed and the standard PCA, the first two principal components accounted for more than 50% of the variance in life-history traits across amniote clades (Figures S8 and S10).

To further validate the findings from the phylogenetic path analysis (PPA) under the Ornstein–Uhlenbeck (OU) model presented in the main text, we repeated the analysis using Pagel's λ model, which was the second-best supported evolutionary framework (Figures S2-S4). The results of these sensitivity analyses are provided in the Supplementary Materials section 2 (Figure S12; Table S7).

### Section 6: Multivariate Ornstein–Uhlenbeck modelling

*(1) Overview of the multivariate OU model*

The multivariate Ornstein–Uhlenbeck (OU) process extends the univariate OU process (Butler & King 2004; Hansen 1997), which describes adaptive evolution of a trait toward an optimal value. The univariate OU process is parameterized by two key components: the selection strength ($\alpha$) governing the rate of adaptation, and the optimum trait value ($\theta$). In its multivariate form, this framework models the joint evolution of a $k$-dimensional suite of traits, $\vec{y}(t)$, over a period of time through the following stochastic differential equation:

$$\mathrm{d}\vec{y}(t) = -\boldsymbol{A}\big(\vec{y}(t) - \vec{\theta}\big)\mathrm{d}t + \Sigma_{yy}\mathrm{d}\vec{W}(t), \qquad (1)$$

where $\vec{y}(t)$ is the vector of trait values at time $t$; $\boldsymbol{A}$ is the matrix counterpart of the selection strength parameter $\alpha$, represents rates of adaptation of trait values towards optimum values; $\vec{\theta}$ is the vector representing the optimum values for these traits; $\vec{W}(t)$ is a $k$-dimensional standard Brownian motion. The diffusion matrix $\Sigma_{yy}$ scales the resulting stochastic perturbations, which represent non-directed changes in trait values attributable to unconsidered selective factors, environmental fluctuations, or evolutionary constraints (Bartoszek *et al.* 2012, 2023, 2024; Clavel *et al.* 2015).

To fully understand the coevolutionary dynamics of these traits, it is essential to examine both the $\boldsymbol{A}$ matrix directly, and to study its eigenvalues and eigenvectors (Bartoszek *et al.* 2023, 2024). The diagonal entries of $\boldsymbol{A}$ ($\boldsymbol{A}_{ii}$) represent the rate at which trait $i$ returns to its own optimum $\theta_i$. The off-diagonal entries ($\boldsymbol{A}_{ij}$) quantify coevolutionary coupling between traits, indicating how a deviation in trait $j$ pulls trait $i$ toward ($\boldsymbol{A}_{ij} > 0$) or away from ($\boldsymbol{A}_{ij} < 0$) its optimum $\theta_i$ (Figure M1a). Therefore, a positive $\boldsymbol{A}_{ij}$ suggests a trade-up between trait $j$ and $i$, whereas a negative $\boldsymbol{A}_{ij}$ can be interpreted as a trade-off.

Since $\boldsymbol{A}$ is the coefficient matrix in the OU stochastic differential equation (Eq. 1), each entry of this matrix captures the instantaneous rate of adaption/coevolution. In contrast, the eigenvalues ($\lambda$) of $\boldsymbol{A}$ describe the relative long-term rates at which linear combinations of traits evolve toward their optima along the independent directions in trait space defined by the corresponding eigenvectors ($\vec{e}$) of $\boldsymbol{A}$ (Bartoszek *et al.* 2023, 2024). Each eigenvector represents an independent evolutionary mode (direction) in trait space, with larger eigenvalues indicating faster evolution along those trajectories. To facilitate interpretation, we also reported the evolutionary half-life ($t_{1/2} = \frac{ln2}{\lambda}$) for each eigenvector. This metric represents the time required for an eigenvector to evolve halfway from its ancestral state to the optimum and serves as a measure of phylogenetic signal (Hansen & Bartoszek 2012). For comparative clarity, we also report this half-life as a relative percentage of total tree height.

*(2) Initial model fitting to distinguish the best model defined in PPA*

We implemented all model fitting using the R package "*mvSLOUCH*" (2.7.6; Bartoszek *et al.* 2024). Our first step was to compare the relative likelihood of the 32 previously defined DAGs (Figure S9), which represent hypotheses for the relationships among five life-history traits (OS to CF). This step assessed whether the direct and indirect associations identified by PPA were robust within an alternative multivariate OU modelling framework.

In mvSLOUCH, the structure of each candidate DAG is encoded in the $\boldsymbol{A}$ matrix through parameter constraints (Figure M1b): (i) If a DAG specified no direct link between two nodes (traits), the corresponding entry in the $\boldsymbol{A}$ matrix was fixed at 0. (ii) If a DAG specified a direct link between two nodes, the corresponding off-diagonal entry in the $\boldsymbol{A}$ matrix was set to 'NA', allowing it to be freely estimated. (iii) A direct link could be further specified by a hypothesized direction of effect. In these cases, the corresponding entry was constrained to be either '$+$' (positive) or '$-$' (negative), forcing the model to fit a specific type of coevolution.

For this initial model comparison step, our aim was to distinguish the direct and indirect associations among traits. Therefore, all diagonal entries of $\boldsymbol{A}$ were constrained to be positive, ensuring each trait approaches its own optimum, while all non-zero off-diagonal entries were left as 'NA' (free to fit), allowing the data to determine the sign and strength of each direct coevolutionary coupling without prior assumption (Figure M1b).

To facilitate numerical optimization, we scaled all phylogenetic branch lengths by the total tree height, converting them to proportions of one. Model comparison was based on the Akaike information criterion corrected for sample size (AICc; Hurvich & Tsai 1989), which has been shown to outperform other criteria (e.g., AIC, BIC) for differentiating among various OU model setups (Bartoszek *et al.* 2024). Other criteria are reported only for reference (Table S8).

To ensure robust parameter estimates and confident model selection, each model configuration was run 500 times with different starting points, obtained from preliminary fits with $\boldsymbol{A}$ set to "*DecomposablePositive*". The run with the smallest AICc value was selected as the best parameter estimate for that model.

We also tested whether the traits evolved with correlated or independent stochastic perturbations by fitting each model with two parameterizations of the $\Sigma_{yy}$ diffusion matrix: 'diagonal' (independent perturbations) and 'upper triangular' (correlated perturbations). Across all candidate models, the 'diagonal' parameterization was consistently favoured by a smaller AICc, indicating that stochastic perturbations act independently on each trait (Figure 4a). Consequently, we only analysed models with ‘diagonal’ $\Sigma_{yy}$.

For each animal group, the best-supported candidate model was identified as the one with the smallest AICc value across all runs of all models (Figure M1b). The best models were retained for further parameterization in the subsequent step.

*(3) Testing causal structures and selecting the best directional model*

A subsequent step was performed to infer the most likely direction of coevolutionary coupling within the best-supported model identified in the previous step for each animal group. The asymmetry of the $\boldsymbol{A}$ matrix ($\boldsymbol{A}_{ij} \neq \boldsymbol{A}_{ji}$), provides a framework for this type of causal inference, as each possible direction of an arrow (e.g., Trait A → Trait B vs. Trait B → Trait A) represents a distinct biological hypothesis about which trait exerts evolutionary influence on the other (Reitan *et al.* 2012). To address this, we treated the previously selected best DAG as a skeleton, and systematically tested all possible combinations of directions for its arrows (see Figure M1c for examples), with each combination defining a distinct causal model. Because the best-supported model from our initial step was a subset of the best model identified in the PPA (Figure 4b; Table S8), we constrained the signs (positive or negative) of the non-zero off-diagonal entries in $\boldsymbol{A}$ to match the fitted coefficients from the corresponding PPA model. This incorporated the prior evidence on the nature of the relationships while testing their direction. Model comparison based on AICc was then used to select the specific causal structure (i.e., the arrangement of arrow directions) that was best supported by the data. To ensure robust parameter estimates for this set of candidate models, each was run 100 times from different starting points, and the run with the lowest AICc was reported as the final best model for each animal group. Our conclusions were therefore drawn by analysing the $\boldsymbol{A}$ matrix, and the eigenvectors and eigenvalues of the $\boldsymbol{A}$ matrix from the selected final best model for each animal group (Figure 4; Table 1).

*(4) Sensitivity analyses for the mvOU model*

To improve the robustness of the results in our main text, we expanded the taxonomic sampling of birds. We first used a phylogenetic imputation approach implemented in the R package *“Rphylopars”* (version 0.3.10; Goolsby *et al.* 2017) to estimate missing life-history values while accounting for shared evolutionary history. Avian species with empirical data for at least four of the five focal traits were retained, and the remaining missing values were imputed, resulting in a dataset of 643 species (Table S1).

Because most missing values involved offspring size (OS), we conducted an additional sensitivity analysis in which this trait was excluded, retaining only species with complete records for the four remaining life-history traits with the best data coverage. Removing offspring size allowed us to include a larger number of bird species, yielding a dataset of 738 species (Table S1). We repeated the mvOU analyses for both datasets following the same procedures described above. The results of these sensitivity analyses are provided in the Supplementary Materials section 3 (Figure S13).

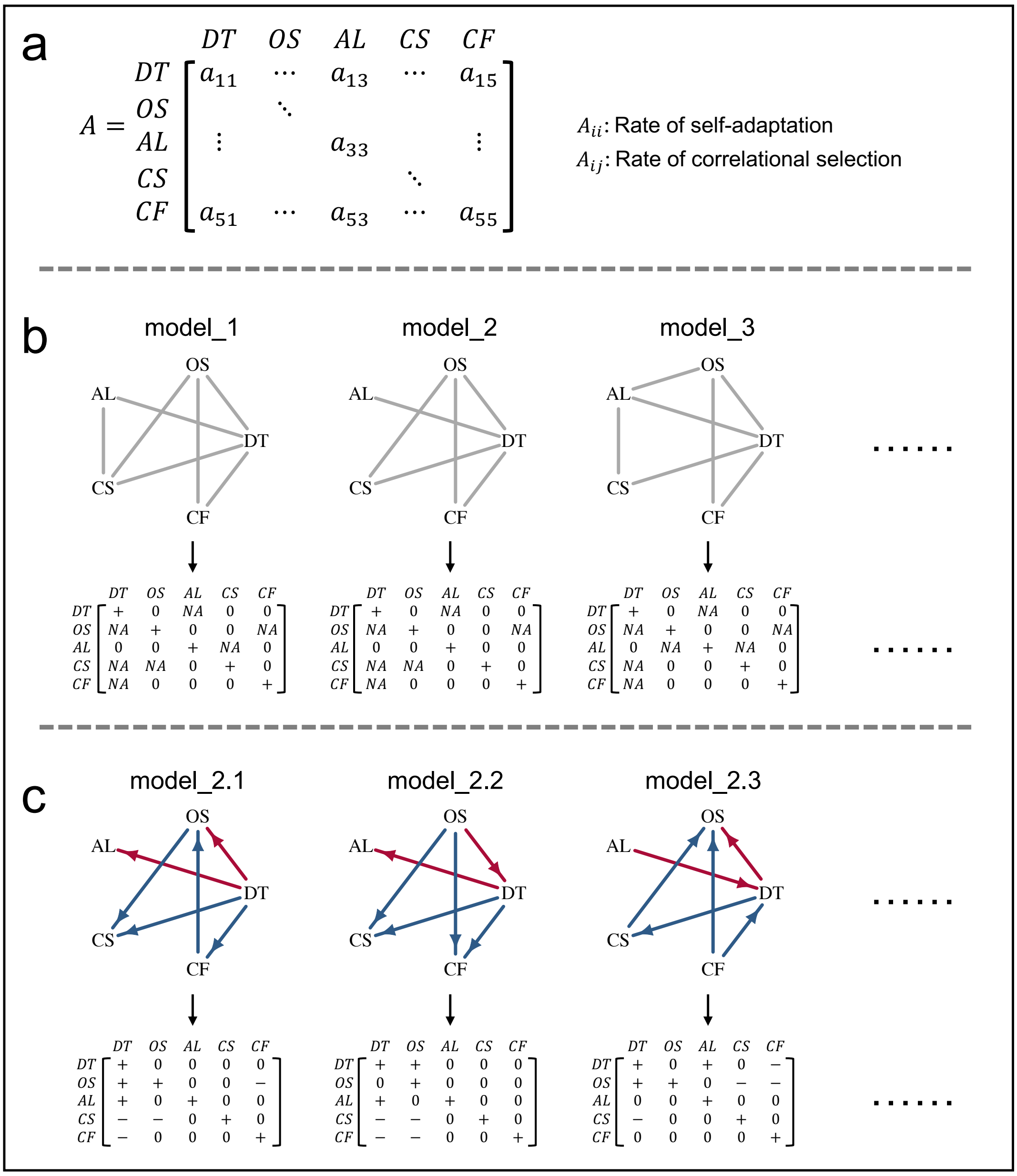


**Figure M1.** Model fitting procedure for multivariate Ornstein–Uhlenbeck (OU) models. (a) The $\boldsymbol{A}$ matrix encodes the instantaneous rate of coevolution among traits (Eq.1). Diagonal elements ($\boldsymbol{A}_{ii}$) represent the rate of self-adaptation. Off-diagonal entries ($\boldsymbol{A}_{ij}$) quantify the correlational selection between traits, representing how a deviation in trait $j$ pulls trait $i$ toward or away from its optimum. (b) Initial model fitting: parameters of the $\boldsymbol{A}$ matrix for each multivariate OU model were constrained based on previously defined DAGs (Figure S9). A missing link between two nodes (e.g., AL ~ OS in model_1) fixed the corresponding entry ($\boldsymbol{A}_{AL,OS}$) at 0. A direct link between two nodes (e.g., DT ~ AL in model_2) allowed the corresponding entry ($\boldsymbol{A}_{DT,AL}$) to be freely estimated (set as 'NA'). All diagonal elements of $\boldsymbol{A}$ were constrained to be positive. (c) Causal structure testing: For the best-supported model from the previous step (e.g., model_2), we tested all possible causal structures (i.e., directions of arrows) on the coevolution among these traits by further constraining elements of $\boldsymbol{A}$. If the value of $\boldsymbol{A}_{ij}$ is positive ('+'), it indicates that trait $j$ pulls trait $i$ toward its optimum, which is represented by a red arrow pointing from $j$ to $i$; conversely, if the value is negative ('−'), it indicates that trait $j$ pulls trait $i$ away from its optimum, which is represented by a blue arrow pointing from $j$ to $i$.
